# A wheat chromosome segment substitution line series supports characterisation and use of progenitor genetic variation

**DOI:** 10.1101/2022.06.18.496684

**Authors:** Richard Horsnell, Fiona J Leigh, Tally IC Wright, Amanda J Burridge, Aleksander Ligeza, Alexandra M. Przewieslik-Allen, Philip Howell, Cristobal Uauy, Keith J. Edwards, Alison R Bentley

## Abstract

Genome-wide introgression and substitution lines have been developed in many plant species, enhancing mapping precision, gene discovery and the identification and exploitation of variation from wild relatives. Created over multiple generations of crossing and/or backcrossing accompanied by marker-assisted selection, the resulting introgression lines are a fixed genetic resource. In this study we report the development of spring wheat chromosome segment substitution lines generated to systematically capture genetic variation from tetraploid (*Triticum turgidum* ssp *dicoccoides*) and diploid (*Aegilops tauschii*) progenitor species. Generated in a common genetic background over four generations of backcrossing, the material is a base resource for the mapping and characterisation of wheat progenitor variation. To facilitate further exploitation the final population was genetically characterised using a high-density genotyping array and a range of agronomic and grain traits assessed to demonstrate the the potential use of the populations for trait localisation in wheat.

## Introduction

Hexaploid wheat (*Triticum aestivum*) arose through natural hybridization between the tetraploid grass species *T. turgidum* and the diploid species *Aegilops tauschii* (Fu & Somers, 2009). The infrequency of hybridisation resulted in limited genetic diversity (Cox, 1997) and progenitor species are a valuable source of variation (Zhang *et al*., 2015). Numerous strategies have been proposed and deployed to capture this for application in genetic studies and wheat breeding (Leigh *et al*., 2022).

A wealth of genomic and germplasm resources is now available for functional genetic characterisation in polyploid wheat (Adamski *et al*., 2020). This includes mapping and gene discovery resources for wild relatives (e.g. stable wild wheat introgression lines developed by Grewal *et al*., 2020). These build on early cytogenetic resources including monosomic and nullisomic chromosome substitution lines series (Sears, 1954; Unrau *et al*., 1956) which supported the foundations of wheat genetics (Kuspira and Unrau, 1957; Law, 1966). The original monosomics took over 25 years to develop (Berke *et al*., 1992) and were used to characterise a range of traits (Zemetra *et al*., 1986) as well as to create reciprocal series of chromosome substitution lines, allowing genetic trait mapping (Berke *et al*., 1992).

Substitution line series serve many purposes. Law (1966) described their benefits as three-fold: (1) localisation of genetic effects enabling targeted use in breeding, (2) revealing genetic trait architecture and relating it to patterns of descent and (3) validating predictions of magnitude of genetic effects. Their direct use in breeding has been demonstrated as base populations for trait mapping and line selection (Eshed *et al*., 1992), identification (Basava, 2019) and transfer of wild species variation (Eshed & Zamir, 1994; Nie *et al*., 2015) and pyramiding of target regions or introgression segments (Ali *et al*., 2010). In understanding trait architecture, they support precise QTL mapping (Basava, 2019) and fine mapping (Fulop *et al*., 2016), gene discovery and cloning (Eshed & Zamir, 1994) along with marker development (Qiao *et al*., 2016). Substitution lines typically show little variation for agronomic (growth and development) traits and can be used to confirm predicted magnitude of genetic effect, thereby increasing homogeneity between experiments (Keurentjes *et al*., 2007) as well as eliminating the interference of genetic background (Zhai *et al*., 2016). They are permanent populations (Zhai *et al*., 2016) that can be maintained as immortal seed stocks allowing assessment across environments and traits (Keurentjes *et al*., 2007), thereby increasing statistical power to detect small effects QTLs and accurately determine the magnitude of genotype x environment interactions (Rae *et al*, 1999; Keurentjes *et al*., 2007).

Chromosome segment substitution lines (CSSLs) are a form of substitution series and capture genome-wide diversity from a donor species in a fixed genetic background. First described in tomato as a series of interspecific introgression lines capturing small, overlapping chromosome segments of the wild species *Lycopersicum pennellii* into cultivated tomato (*L. esculentum*; Eshed *et al*., 1992; Eshed & Zamir, 1994) the resulting lines were used to map quantitative traits including yield (Eshed & Zamir, 1995; Fridman *et al*., 2004) and leaf characters (Holtan & Hake, 2003). The introgression lines have been subsequently used to detect thousands of QTLs for adaptation, morphological characters, yield, metabolism and fruit quality traits (reviewed in Lippman *et al*., 2007) and cellular features underlying leaf traits (Chitwood *et al*., 2013).

Further application has been demonstrated in the model species *Arabidopsis thaliana* (Keurentjes *et al.,* 2007; Fletcher *et al*., 2013) as well as in several cultivated crops including rice (reviewed by Ali *et al*., 2010), brassica (Howell *et al*., 1996; Ramsay *et al*., 1996; Rae *et al*., 1999; Li *et al*., 2015), peanut (Fonceka *et al*., 2012), durum wheat (Blanco *et al*., 2006) and pearl millet (Kumari *et al*., 2014; Basava *et al*., 2019). A recent review of resources by Balakrishnan *et al*. (2019) detailed populations available over sixteen cultivated crop species. Despite the importance of wheat in global food security and its limited diversity due to evolutionary history (Stebbins, 1950; Gross & Olsen, 2010) there are few substitution resources available beyond the foundation monosomic and nullisomic series. In hexaploid wheat Zemetra *et al*. (1986) used the Nebraska reciprocal CSSL series to map heading date. More recently Gu *et al*. (2015) developed four sets of hexaploid introgression lines capturing the endemic Chinese wheat subspecies *T. aestivum yunnanense*, *tibetanum* and *petropavlovskyi* along with a synthetic hexaploid wheat. The lines were used to map QTLs for height, spike length and grain number per spike and several lines were identified for use in breeding to incorporate favourable introgression segments. In tetraploid wheat, Joppa and Cantrell (1990) created a ‘Langdon’(durum)-*T. dicoccoides* substitution series via crossing between a high grain protein content (GPC) *T. dicoccoides* donor and a series of ‘Langdon’ D-genome disomic substitution lines. This allowed the identification of lines with superior GPC, subsequently leading to its mapping on chromosome 6B (*QGpc.ndsu-6Bb*; Joppa *et al*., 1997). Further refinement of mapping to the 6B short arm (Olmos *et al*., 2003) provided the basis for the cloning of the *GRAIN PROTEIN CONTENT-B1* (*GPC-B1*) gene (Uauy *et al*., 2006a; Uauy *et al*., 2006b).

This demonstrates the utility of precision introgression resources for trait mapping and gene cloning. In durum wheat, Blanco *et al*. (2006) also developed a set of 92 backcross introgression lines using *T. turgidum* ssp. *dicoccoides*. Developed to capture high GPC from the wild species, several generations of backcrossing allowed extraction of a single favourable line. This was used to derive a population with the original recurrent parent and to extract high and low protein bulks from the derived F3 population for marker discovery and QTL detection. Selection of CSSLs within a series aims to capture segments supporting genetic dissection and that optimise donor coverage. Balakrishnan *et al*. (2019) reviewed the breadth and characteristics of existing CSSL resources reporting that sets were composed of 35-200 lines, typically capturing 90% of donor genome. Tian *et al*. (2006) reported 67% coverage for wild to cultivated rice compared to near complete (99%) coverage in other rice population (Jie *et al*., 2006). In supporting genetic dissection, substitution line phenotypic effects that differ between an introgression line and the recurrent parent are ascribed to the substituted region (Rae *et al*., 1999). To optimise this Wu *et al*. (2006) proposed an additive-dominance model for accurately determining and ascribing substituted segment effects.

In this study we describe the development and characterization of a CSSL series capturing the genomes of hexaploid wheat’s primary progenitor species: tetraploid wild emmer (*T. turgidum ssp. dicoccoides;* AB genomes) and diploid goat grass (*Ae. tauschii;* D genome). We genetically characterize the resource using a high-density marker platform to provide accurate resolution of introgression boundaries. We demonstrate that the series can be used to associate trait variation from progenitor species to specific genetic regions. The series is publicly available to provide characterised introgression lines (and associated markers) to support further trait research and progression into breeding.

## Materials and Methods

### Selection of parental material

Two independent backcross-derived CSSL populations were generated to capture the genomes of the progenitor species *T. turgidum ssp. dicoccoides* (denoted NIAB_AB) and *Ae. tauschii* (denoted NIAB_D). The hexaploid spring wheat ‘Paragon’ was used as the recurrent parent for both populations as it has been widely used for genetic studies and the creation of genetic stocks. For the NIAB_AB population, the Israeli wild emmer wheat accession TTD-140 was selected as the donor parent. For the NIAB_D population, the Armenian *Ae. tauschii* accession ENT-336 was selected and the synthetic hexaploid line NIAB-SHW041 (created via resynthesis from a Hoh-501 (AB) x ENT-336 (D) cross) used as the introgression donor.

Comparative genetic analysis of ‘Paragon’, TTD-140 and NIAB-SHW041 used single nucleotide polymorphism (SNP) marker data generated using the Axiom® Wheat Breeder’s Genotyping Array (Allen *et al.,* 2017). Two datasets were used, the first with A and B genome mapped markers and ‘Paragon’, 24 *T. dicoccoides* (including TTD-140), 12 *T. dicoccum*, 12 *T. durum* accessions and the second using only D genome markers and ‘Paragon’ and 51 primary synthetic hexaploid wheats (including NIAB-SHW041). Markers were thinned using pairwise comparisons between SNPs using a Pearson correlation test and a single marker was removed in comparisons that yielded an absolute value of the correlation coefficient (*r*) greater than 0.75. The final marker number in the A/B and D genome datasets 2,748 and 217, respectively. The relationships between individuals in each set were compared using principal coordinate analysis (PCoA) of Euclidean genetic distance matrices computed from the marker data, implemented in the R package ape (v5.3, Paradis & Schliep 2018).

### Construction of populations

For NIAB_AB, pollen from the wild emmer wheat donor TTD-140 was used to fertilise ‘Paragon’ to create a pentaploid F1 generation. Six F1 plants were used as the maternal parents in the creation of the BC1 generation, where ‘Paragon’ was used to pollinate F1 plants, continuing to BC4. For NIAB_D the donor ENT-336 was first crossed to the durum wheat Hoh- 501 to produce a fertile hexaploid synthetic wheat (NIAB-SHW041). The synthetic wheat was crossed to ‘Paragon’ and the F1 progeny then backcrossed four times to ‘Paragon’ to derive BC4 plants. At each BC generation genomic DNA was extracted from 2-week-old seedlings of all progenies using a modified Tanksley extraction protocol (Fulton *et al*., 1995). Marker-assisted selection (MAS) was used to select individuals carrying heterozygous introgressions in the F1 to BC4 generations and homozygous introgressions in the BC4F2 generation. A schematic of the crossing scheme is given in **Supplementary Figure S1**. The number of lines generated, genotyped, and advanced at each generation is summarised in **Supplementary Table S1**.

MAS at each BC generation used co-dominant KASP™ markers developed to cover the TTD-140 and ENT-336 genomes and to allow for differentiation from ‘Paragon’. In addition to foreground selection, evenly spaced markers on the non-target genomes were used to allow background selection against the donor. At BC4F4 additional genome-wide SNP genotyping (using the Axiom® Wheat Breeder’s Genotyping Array; Allen *et al*., 2017) was used to improve the accuracy of final selection.

For the NIAB_AB population, 191 KASP™ markers (98 A genome; 93 B genome) distributed at approximately 20-30 cM intervals were selected to screen the BC generations (**Supplementary Table S2**). An additonal set of 50 co-dominant D-genome mapped markers (1-2 per chromosome arm) was used to facilitate selection against tetraploids and nullisomic aneuploids and amplified a product in D-genome diploids and hexaploids, but not in AB- genome tetraploids, and gave expected results across nulli-tetrasomic series (**Supplementary Table S3**).

Based on graphical genotypes derived from KASP™ marker positions, subsets of BC1 lines were manually created. The BC1 lines were selected to carry multiple but complementary heterozygous introgressions across the TTD-140 genome. Lines that failed to amplify at loci targeted by D genome specific KASP™ markers were discarded. For each subsequent cycle of backcrossing, the same selection criteria were applied. A summary of the number of individuals created, genotyped, and selected at each crossing generation is given in **Supplementary Table S1**. From each of the 138 selected BC4F1 individuals, 8 BC4F2 seeds were grown and genotyped (using the above KASP^TM^ marker set; **Supplementary Table S2**) and one individual was selected based on homozygosity for the marker at the target introgression but minimising background introgressions (using markers in **Supplementary Table S3**). Eight BC4F3 seeds from each selected homozygous BC4F2 line were subsequently sown and genotyped with the KASP™ marker set to identify the target introgression and to ensure progression of homozygous lines.

At the BC4F4 generation, the lines were genotyped using the Axiom® Wheat Breeder’s Genotyping Array (Allen *et al*., 2017). To derive the final NIAB_AB population we used 1,444 A-genome and 1,707 B-genome SNPs which were verified to share the same genetic and physical genome chromosome. Genetic positions were taken from the existing consensus map (Allen *et al*., 2017) and physical positions determined using the Basic Local Alignment Search Tool (BLAST+) command-line applications on the RefSeq v1.0 genome assembly with default parameters (Camacho *et al*., 2009, International Wheat Genome Sequencing Consortium (IWGSC) 2018). Only markers with a single A or B genome physical hit were used. Introgression selections were made using physical positions and graphical visualisation of allele variation different from ‘Paragon’ using Flapjack (v-1.18.06.29, Milne *et al*., 2010). Introgressions were visually selected on: (a) coverage across the target genome chromosome, (b) homozygosity within each introgression and (c) minimal off-target introgressions.

For the NIAB_D population, 60 D-genome KASP™ markers were selected based on distribution across the D-genome, co-dominance and polymorphism between the recurrent parent ‘Paragon’, the tetraploid component of NIAB-SHW041, Hoh-501 and the *Ae. tauschii* donor ENT-336 (**Supplementary Table S4**). In addition, 66 A- (33) and B-genome (33) markers were used for background selection against Hoh-501 (**Supplementary Table S5**). The CSSLs were developed using MAS as described for the NIAB_AB population but with ‘Paragon’ used as the maternal parent, as summarized in **Supplementary Table S1**.

At the BC4F4 generation, lines were genotyped using the Axiom® Wheat Breeder’s Genotyping Array. Using the same approach as for the NIAB_AB population, 644 SNPs which shared a genetic and physical D genome chromosome (where markers had only a single physical D genome hit) were used for the final line selection. Introgression line selections were based on coverage across each chromosome, maximising homozygosity and minimising off-target introgressions. Seed of both the NIAB_AB and NIAB_D populations is available from the Germplasm Resources Unit at the John Innes Centre (entries WCSSL0001 to WCSSL0112; available via https://www.seedstor.ac.uk/) and all genotypic data is available from www.cerealsdb.uk.net/cerealgenomics/CerealsDB/array_info.php.

### Phenotypic characterisation

The NIAB_AB population was grown under glasshouse conditions to assess grain yield per plant and grain characteristics. Two or three replicate plants of each line were grown in an incomplete block design, surrounded by a ‘Paragon’ border to minimise edge effects. Each plant was grown in a 1L pot with 16-hour daylength at 22°C and 15°C overnight. At maturity, all plants plus one ear per plant were photographed, and all plant images are available from: https://opendata.earlham.ac.uk/wheat/under_license/toronto/Horsnell,Leigh,Bentley,Wright_ <u>2020_NIAB_CSSL_D_genome_yield_trial_H2020/</u>. Seed from each plant was hand threshed and weighed to estimate grain yield per plant. Grains were analysed using a MARVIN seed analyser (MARViTECH GmbH) to obtain grain dimensions and thousand grain weight (TGW).

To assess phenotypic variance in the NIAB_D CSSLs, three replicates of each line along with six replicates of the recurrent parent ‘Paragon’ were sown in a randomised yield (2x6 m harvested area) trial in Hinxton, Cambridge in 2020. The trial was laid out in an incomplete block design, with 6 plots per block and three complete replicates. Traits assessed in the field included: grain yield (kg/plot at 85% dry matter content), early ground coverage using normalised difference vegetation index (NDVI^early^; an average of two measurements taken between 11:00 and 14:00 GMT on the 15^th^ and 21^st^ of April 2020 at 2m above the canopy using a handheld RapidScan CS-45 (Holland Scientific)), plant height (cm from base to tip (including scurs or awns) of three random plants per plot at maturity), flowering time (days to growth stage (GS) 61 (start of anthesis) recorded when 50% of the plot was at GS61; Zadoks *et al*., 1974). All field trial data is available from Grassroots (Bian *et al*., 2017): https://grassroots.tools/fieldtrial/study/61faaf25c68884365e7bcc34. Images of each plot were taken to assess glaucosity using a RGB camera (Olympus TG4) at a height of 320cm from the ground. Images were cropped to eliminate edge distortion with the final 8 Mpix image covering 85% of the plot. Post-processed images for each plot are available here: https://opendata.earlham.ac.uk/wheat/under_license/toronto/Horsnell,Leigh,Bentley,Wright_ <u>2020_NIAB_CSSL_D_genome_yield_trial_H2020/</u>. Post-harvest yield component traits were assessed on ten replicate ears per plot (collected at random) including ear length (cm from base to tip of the ear, not including scurs or awns) and spikelet number. Each of the 10 ears was weighed to determine ear weight, threshed individually and grains analysed using a MARVIN seed analyser (MARViTECH GmbH) to determine seed number, thousand grain weight (TGW) and grain dimensions (length, width).

### Phenotypic analysis

For both the NIAB_AB glasshouse and NIAB_D field trial, analysis was conducted in R v4.0.3 (R Core Team, 2020). Each trial included the final CSSLs selected to form the graphical genotype as well as a proportion of CSSLs that were not selected in the final population but carried introgressions. The NIAB_AB glasshouse trial analysis included 146 replicated CSSL individuals. In the NIAB_D field trial analysis there were 33 replicated CSSLs. For each trait, variance between replicates and population distributions were plotted and visually inspected to remove outliers. In the NIAB_D field trial, 10 ears were harvested per plot as technical replicates for measurements on ear and seed characteristics. The ears were measured separately and variance within a plot was then inspected to aid in outlier removal. The technical replicates were then averaged to form plot means for the analysis.

Mixed linear effect models were constructed with restricted maximum likelihood (REML) using the R packages lme4 (Bates *et al.,* 2015) and lmerTest (Kuznetsova *et al.,* 2017). Genotype was treated as a fixed factor and the trial design components including ‘replicate’ and ‘block’ were tested within each model as random factors. Due to visual patterns in model residuals, in addition to the direction of phenotypic assessment and application of agronomic inputs, the trial layout components ‘column’ in the NIAB_D and ‘row’ in the NIAB_AB glasshouse trial were included as random factors. Final model selection was based on Akaike Information Criterion (AIC) and (exact) restricted likelihood ratio tests implemented through the R package RLRsim (Scheipl *et al.,* 2008). In each model, the significance of the genotypic effect was estimated using ANOVA with Satterthwaite’s degrees of freedom method, implemented through lmerTest. Generalised heritability (*H^2^*) was estimated using genotypic best linear unbiased predictors (BLUPs) and the Cullis *et al*. (2006) model, using the R script developed by Schmidt *et al*. (2019). Ranked bar plots, including only the final selected CSSLs, were created for grain yield (NIAB_D trial), seed width (NIAB_D trial) and seed length (NIAB_AB trial). The package emmeans (Lenth, 2020) was used to complete pairwise comparisons of all BLUEs for these traits and a false discovery rate (FDR) correction was used to adjust *P* values for multiple testing. Finally, the package predictmeans (Luo *et al.,* 2020) was used to inspect the residuals from final models and to extract the BLUEs for examining variation across all traits.

### Introgression-trait association

Categorical traits were assessed in each CSSL population to demonstrate the use of the resource for detecting introgression-trait association. In NIAB_AB this used a major known locus linked to awn presence and in NIAB_D to a putative marker association with flag leaf glaucousness. Awn presence was scored in the NIAB_AB glasshouse trial and graphical genotypes were inspected to detect introgressions from the awned donor TTD-140 overlapping with regions where major awn inhibitors have previously been mapped. Huang *et al*. (2020) characterised and identified the *B1* locus as a C2H2 zinc finger encoding gene that corresponds to a coding region on chromosome 5A (698,528,636 to 698,529,001bp; IWGSC 2018). This region was taken as the physical location of the gene for the introgression-trait association.

To demonstrate that CSSLs can also be used to dissect the genetic basis of a less well characterised association, ear, stem, and leaf glaucousness was scored using visual assessment and RGB images in the NIAB_D field trial. This identified a subset of individuals that shared the non-glaucous phenotype, and these were assessed for common introgression regions. A previous study (Würschum *et al.,* 2020a) detected a putative marker-trait association linked to flag leaf glaucousness mapped to 2D (∼2.9 Mb) and the ability of the series to validate this interval was assessed via introgression-trait association. Off target regions were also inspected to confirm that the subset shared no other common introgressions. Once each introgression- trait association was identified, schematics using graphical genotype selections were formed for each association and a SNP cluster plot was extracted from Axiom Analysis Suite (Thermo Fisher Scientific) that confirmed the CSSL region segregating within the introgressions of interest. For both populations, markers that were excluded or dropped during the QC were inspected visually to improve resolution of introgression boundaries.

## Results

### Genetic relationships between parents

The genetic relationships between 48 tetraploid wheat accessions and 51 D genome donors (as novel SHWs) were compared based on Euclidean genetic distance and plotted through PCoA (**Supplementary Figure S2).** TTD-140, the tetraploid donor of the NIAB_AB population clustered with most of the *T. dicoccoides* lines, though the *T. dicoccoides* accessions were shown to contain significant diverse (**Supplementary Figure S2A)**. The main cluster of *T. dicoccoides* was separate from the cultivated wheats by PCoA1, which explained 12% of the variation. The domesticated wheats, including accessions of *T. durum*, *T. dicoccum* and ‘Paragon’, all clustered above zero on PCoA1. These cultivated groups were separated by the second axis which explained 9% of the variation.

The synthetic donor used for the NIAB_D population (NIAB-SHW041) clustered closely with recurrent parent ‘Paragon’ on the first PCoA axis (**Supplementary Figure S2B**), where there was a distinct group of primary synthetics that clustered separately from the rest of the material towards the right of the plot (slightly above 5 on PCoA1). PCoA1 accounted for 20% of the variation compared to 12% PCoA2. However, within the lineage shared by NIAB-SHW041 and ‘Paragon’, the two lines do not appear closely related, which is supported by their separation on PCoA2 in **Supplementary Figure S2B**.

### Selection of introgression lines based on graphical genotypes

Final lines were selected based on graphical genotypes from the BC4F4 (**Supplementary Table S1**). Two different sets of SNP markers were used for line selection in NIAB_AB: 1,444 markers with physical locations on the A genome and 1,707 markers with physical positions on the B genome. From these data sets, 44 genotypes were identified that had A genome introgressions from TTD-140 (**Figure 1A**). Using the B genome data set, 33 genotypes were identified with introgressions covering the B genome (**Figure 1B**). These 77 selections originated from 62 unique NIAB_AB individuals with some genotypes selected for introgressions at multiple loci. For example, NIAB_AB 1A.04 and 5A.05 represent the same unique individual but are included twice to represent distinct target introgressions. Using the NIAB_D population and a D genome marker set of 644 SNPs, 32 genotypes were selected with introgressions from NIAB-SHW041 (**Figure 1D**). These selections originated from 25 unique NIAB_D lines.

**Figure 1.**
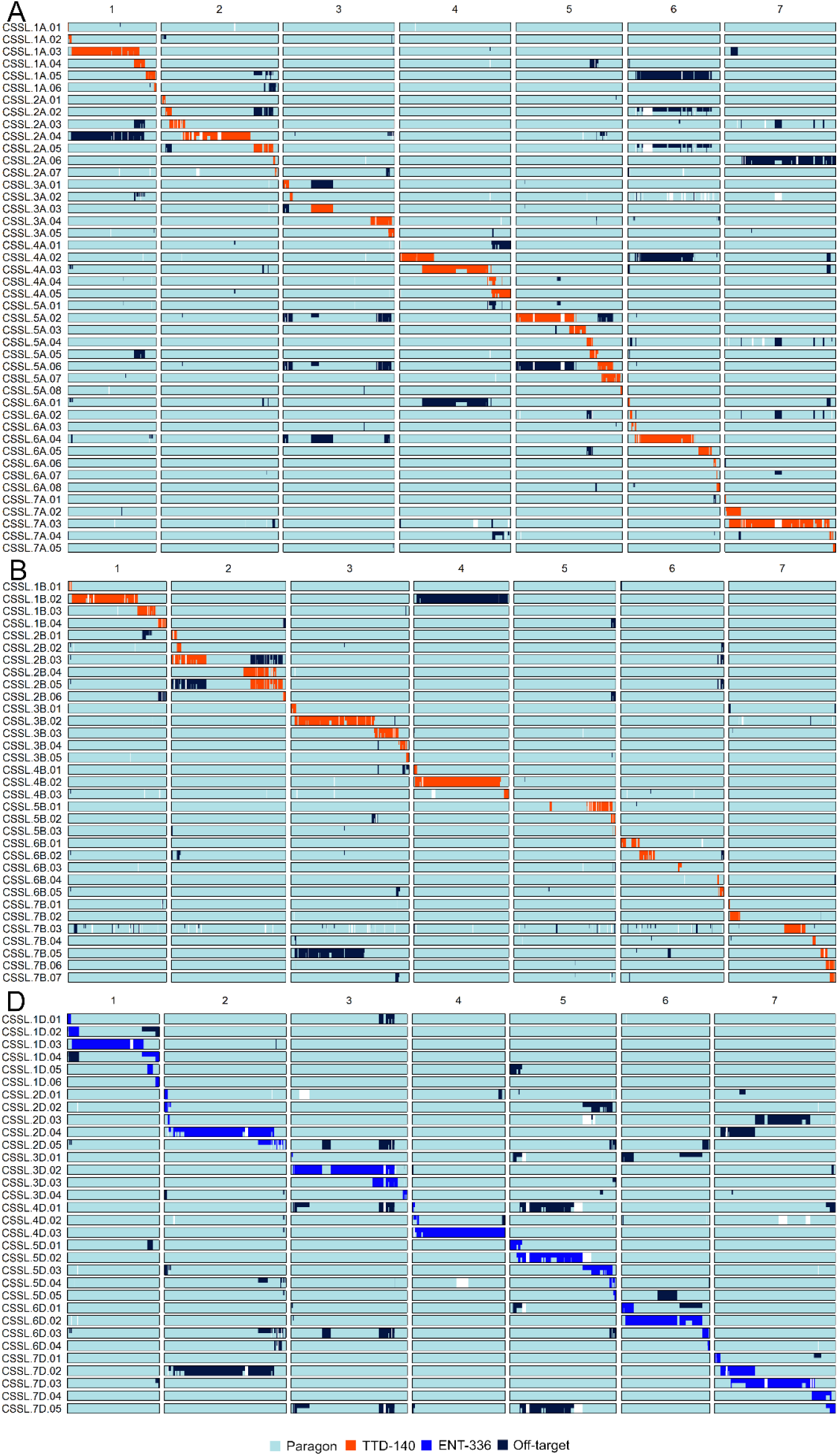
Graphical genotype selections for the A, B and D genome, using the NIAB_AB and NIAB_D populations. A selection of 77 lines was used to form the A and B graphical genotype, originating from 62 unique NIAB_AB CSSLs. The light blue colour represents a ‘Paragon’ like genotype, whereas orange represents a TTD- 140 introgression. Off-target introgressions are shown by navy blue. For the D graphical genotype, 32 lines were selected using 25 unique NIAB_D CSSLs. The medium blue represents an introgression from the NIAB-SHW041 donor, which was formed using the *Ae. tauschii* ENT-336. The chromosomes are scaled on physical position (IWGSC 2018) and the length of each chromosome is determined by marker coverage. White spaces show missing data and heterozygous regions are represented by divided colours. The figure was created using the R package SelectionTools (v-19.1, www.uni-giessen.de).

Individuals were typically selected for a single introgression and genotypes with multiple introgressions were selected against. However, the majority of BC4F4 individuals carried small off-target introgressions (**Figure 1**). Within each genome, each selected genotype contained on average two introgressions (**Figure 1; Supplementary Table S6**). The average length of the introgression ranged from an average of 102 Mb for the B genome to 129 Mb for the D genome. Off-target introgressions were also present across the different genomes for each selected line (**Supplementary Figures S3-S5)**. Therefore, as an estimate of recurrent parent background recovery, the percentage of markers with a ‘Paragon’ allele was quantified for each genotype, using markers with mapped genetic positions on the consensus map (Allen *et al.,* 2017) and genotypes had 96% average recurrent parent recovery (**Supplementary Table S6**). Several NIAB_AB lines appeared to have large chromosomal introgressions from TTD- 140 across the D genome (**Supplementary Figures S3, S4)**. As the tetraploid donor lacks a D genome, these lines were likely D-genome aneuploids and alternative (not ‘Paragon’) alleles were called for markers in that region. The extent of amplification of homeologous loci is not characterised for the markers described herein, although D genome amplification in tetraploid accessions indicates that for some markers, priming sites are sufficiently conserved across genomes to allow amplification.

### Phenotypic variation

Variation was observed for every trait measured in comparison to the recurrent parent ‘Paragon’. Trait summary statistics from the NIAB_AB glasshouse and NIAB_D field trial are shown in **Table 1**. For grain length in both trials, and plant height in the NIAB_D field trial, the mean of the CSSL lines was on average higher than ‘Paragon’ (**Table 1**). However, most trait means for the CSSLs were lower than the recurrent parent, although individual CSSL lines had higher means than ‘Paragon’ for all traits. In the NIAB_AB glasshouse trial there was a 43.4% increase in grain yield per plant in the highest yielding CSSL line (compared to ‘Paragon’) and in NIAB_D field trial the highest TGW for a CSSL line gave a 32.5% increase over ‘Paragon’. For other traits, such as spikelet number and flowering time in NIAB_D, there were more moderate increases in the maximum CSSL mean compared to ‘Paragon’ (**Table 1).** A significant genotypic effect (*P* < 0.001) was observed for all traits (**Table 1**). In NIAB_D, moderate or high generalised heritability (*H^2^*) was observed. The highest *H^2^*was found for grain length (*H^2^*= 0.91) and flowering time, ear length and seed area also showed high *H^2^* (**Table 1**). Estimates of *H^2^* were lowest for grain yield in the NIAB_AB glasshouse assessment (*H^2^* = 0.39), likely due to measurement on a per plant basis.

**Table 1.** Summary statistics and results from two replicated trials: NIAB_AB glasshouse and NIAB_D field trial. Results are taken from all introgressed lines included in both trials and the recurrent parent ‘Paragon’. The best linear unbiased estimates (BLUEs) are shown for ‘Paragon’ and the CSSLs that had the maximum and minimum BLUE for each trait. The genotypic effect for each trait is shown through ANOVA, using the Satterthwaite’s degrees of freedom method. Generalized heritability (*H^2^*) is shown for every trait. For the NIAB_D field trial, ear and seed characteristics (including seed weight) are shown on a per ear basis, averaged from 10 technical replicates per plot.

| Trait | Paragon | CSSL<br>min | CSSL<br>mean | CSSL<br>max | Num<br>df | Den<br>df | F | P | H<sup>2</sup> |
| --- | --- | --- | --- | --- | --- | --- | --- | --- | --- |
| **NIAB_AB glasshouse trial** |  |  |  |  |  |  |  |  |  |
| Grain yield<br>(g/plant) | 4.38 | 1.06 | 3.74 | 6.28 | 143 | 161.36 | 1.67 | <0.001 | 0.39 |
| Grain length<br>(mm) | 6.32 | 5.83 | 6.53 | 7.14 | 144 | 204.02 | 3.56 | <0.001 | 0.70 |
| **NIAB_D field trial** |  |  |  |  |  |  |  |  |  |
| Grain yield<br>(85% DMC kg/plot) | 6.13 | 4.90 | 5.89 | 6.73 | 33 | 55.85 | 3.07 | <0.001 | 0.68 |
| Flowering time<br>(Days from drilling to GS65) | 189.13 | 183.99 | 189.05 | 191.75 | 33 | 54.33 | 8.34 | <0.001 | 0.88 |
| Plant height<br>(cm) | 97.65 | 92.31 | 97.84 | 104.68 | 33 | 57.62 | 4.17 | <0.001 | 0.76 |
| NDVI<sup>early</sup><br>(index) | 0.64 | 0.52 | 0.59 | 0.67 | 33 | 51.97 | 2.94 | <0.001 | 0.65 |
| Spikelet no.<br>(count) | 22.29 | 20.22 | 21.57 | 22.93 | 33 | 59.68 | 6.51 | <0.001 | 0.83 |
| Ear length<br>(cm) | 10.57 | 9.51 | 10.33 | 11.83 | 33 | 59.10 | 8.76 | <0.001 | 0.88 |
| Ear weight<br>(g) | 2.95 | 2.38 | 2.77 | 3.51 | 33 | 56.52 | 2.81 | <0.001 | 0.64 |
| Seed no.<br>(no. ear<sup>-1</sup>) | 58.62 | 42.81 | 56.43 | 63.64 | 33 | 57.46 | 5.88 | <0.001 | 0.82 |
| Seed weight<br>(g ear<sup>-1</sup>) | 2.39 | 1.91 | 2.21 | 2.89 | 33 | 56.65 | 3.20 | <0.001 | 0.68 |
| TGW<br>(g) | 40.56 | 31.69 | 39.10 | 53.74 | 33 | 59.21 | 7.66 | <0.001 | 0.87 |
| Grain length<br>(mm) | 6.63 | 6.35 | 6.70 | 7.33 | 33 | 64.46 | 11.22 | <0.001 | 0.91 |
| Grain width<br>(mm) | 3.43 | 3.13 | 3.35 | 3.65 | 33 | 59.68 | 5.09 | <0.001 | 0.80 |
| Grain area<br>(mm<sup>2</sup>) | 16.80 | 14.66 | 16.55 | 20.12 | 33 | 60.15 | 8.65 | <0.001 | 0.88 |

In the NIAB_AB glasshouse trial, significant variation was observed for grain length (**Table 1**, **Figure 2A**). The majority of the CSSLs had a higher mean than ‘Paragon’; compared to the recurrent parent there was a 13.0% increase in the CSSL with the maximum grain length (NIAB_AB 5A.02/5A.06). For the NIAB_AB selections, post hoc tests indicated several lines with grain length significantly different to ‘Paragon’ (**Figure 2A**) and variation for a subset of lines is shown in **Figure 2B**, clearly demonstrating visual variation. Grain length showed a positive weak correlation with grain yield per plant (*r* = 0.3, *P* = 0.02; **Figure 2C**). Overall, there was a significant genotypic effect observed for grain yield per plant (**Table 1**), although the post hoc tests did not identify any CSSLs that differed significantly to ‘Paragon’ and measurement on a per plant basis in the glasshouse is the likely cause of the large standard error associated with the BLUEs (**Figure 2D**).

**Figure 2.**
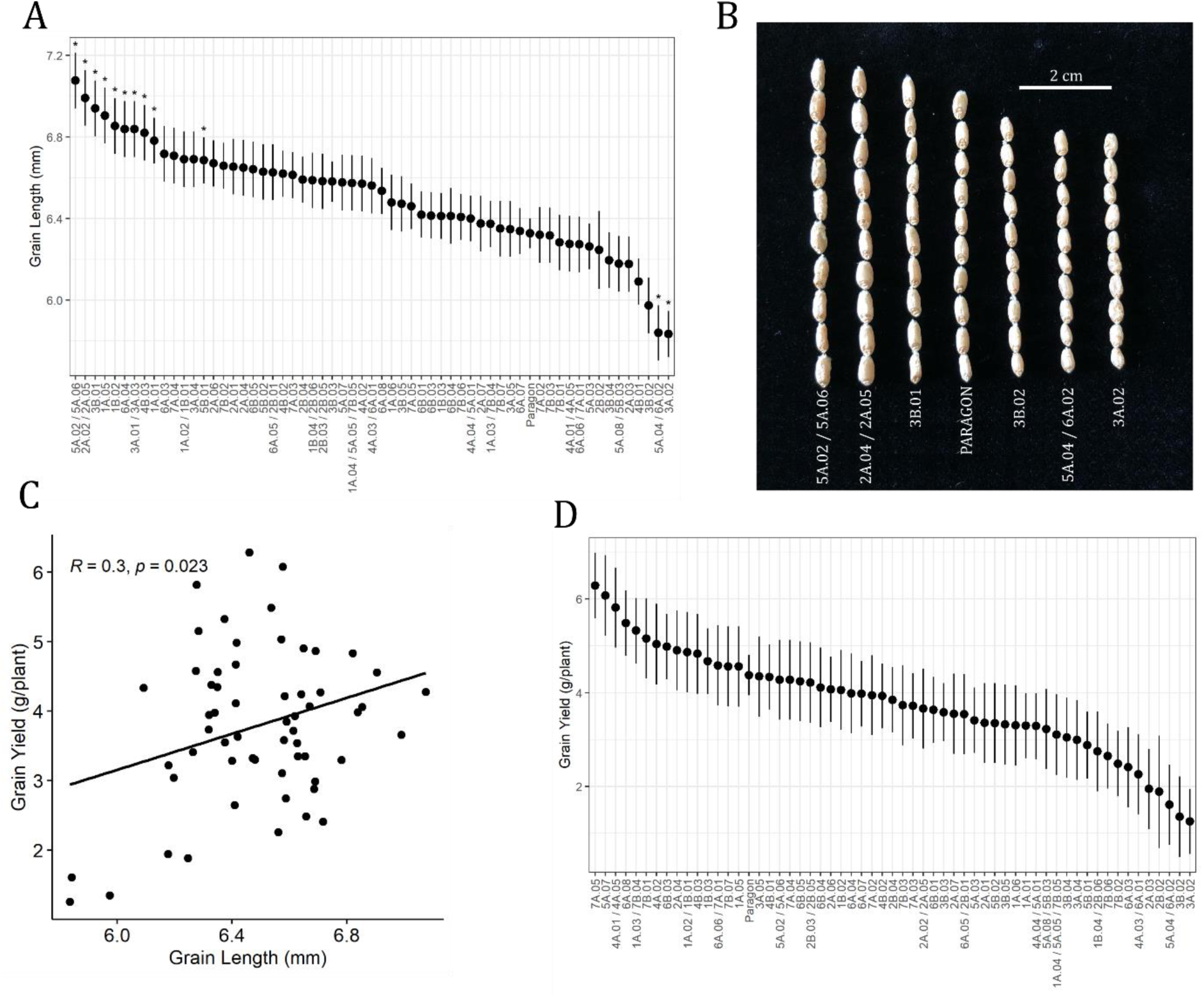
Variation for grain length and yield per plant in the NIAB_AB glasshouse trial: A) Ranked best linear unbiased estimates (BLUEs) for seed length, with the NIAB_AB lines selected for the graphical genotype and the recurrent parent ‘Paragon’ shown. The asterisks above genotypes indicate where CSSLs were significantly different to ‘Paragon’ (*P* < 0.05). B) Variation in seed length observed with CSSLs with the longest and shortest seeds included, and ‘Paragon’ shown for comparison. C) The relationship between grain length and yield, with the Pearson correlation coefficient test statistics overlayed on the plot. D) NIAB_AB CSSLs and ‘Paragon’ ranked BLUEs for grain yield per plant. Error bars in A and D represent standard error of the BLUEs.

Several traits showed a positive correlation with grain yield in the NIAB_D trial (**Figure 3A)**, of which the positive correlation with grain width was the most significant (*r* = 0.61, *P* < 0.001; **Figure 3A**). Several significant negative trade-offs were also observed with flowering time having the most negative correlations with other traits (**Figure 3A)**. Earlier flowering was negatively associated with ear length, TGW and grain length, width, and area. A significant negative relationship was also observed between seed number (per ear) and other grain size related traits (including TGW, seed length and area).

**Figure 3.**
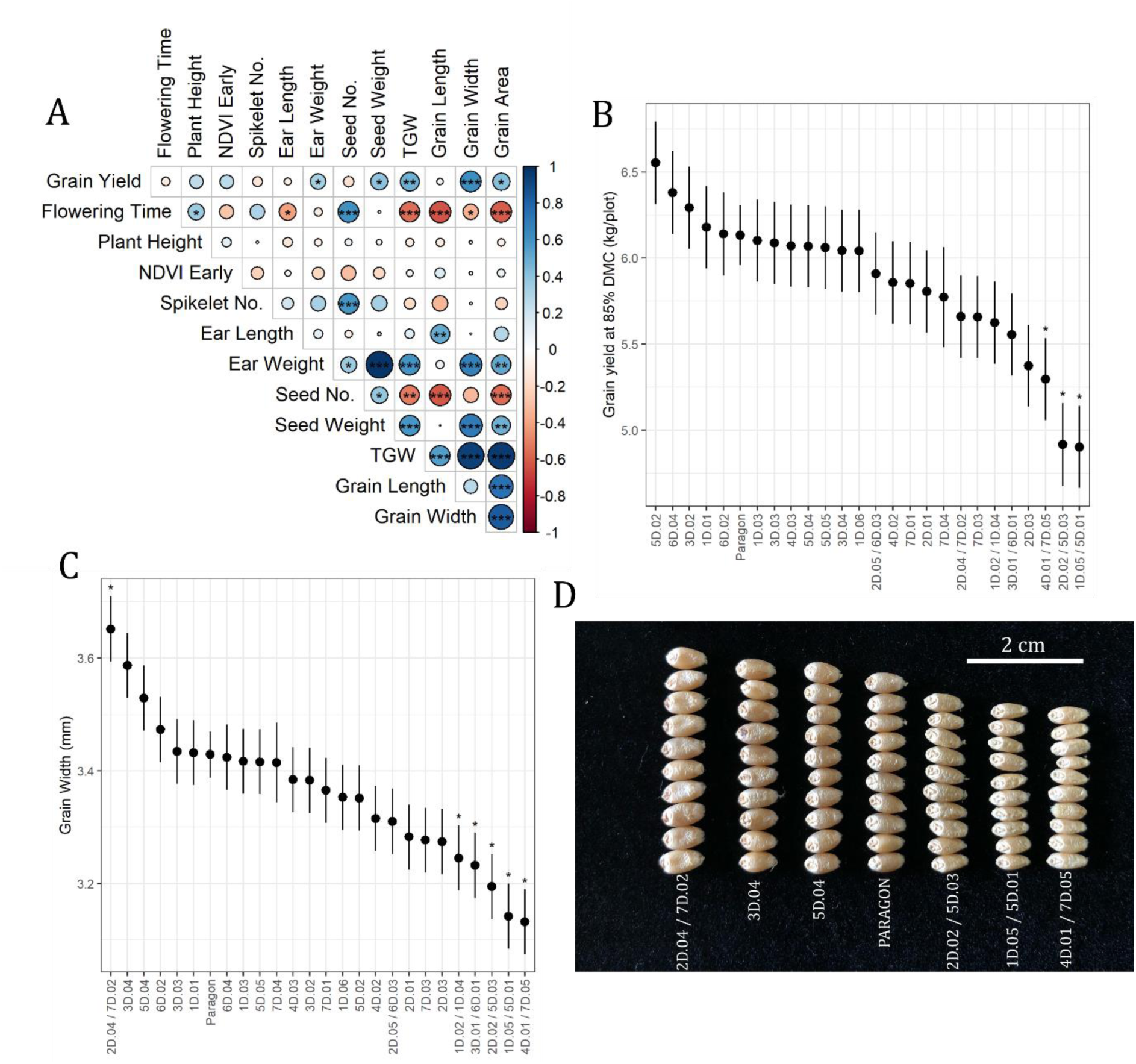
Trait correlations and variation for grain yield and grain width in the NIAB_D field trial: A) a Pearson’s correlation matrix calculated using the best linear unbiased estimates (BLUEs) from models fitted for each trait.

Grain yield was reduced in the majority of selected CSSLs compared to ‘Paragon’, except in five CSSLs (**Figure 3B**). Lines selected for multiple introgressions across the D genome gave some of the lowest yield performance. Post-hoc pairwise comparisons showed that three of these were significantly lower than ‘Paragon’ (4D.01/7D.05, 2D.02/5D.03 and 1D.05/5D.01). None of the CSSLs had significantly higher grain yield than ‘Paragon’. Several NIAB_D CSSLs showed significantly different means for grain width compared to ‘Paragon’: five CSSLs had a lower predicted mean and a single CSSL had a higher mean (2D.04/7D.02, **Figure 3C**). There were clear visual differences in grain width observed across the NIAB_D CSSLs (**Figure 3D**). However, despite wide grain width CSSL.2D.04/7D.02 did not have a higher grain yield than ‘Paragon’ (**Figure 3B**).

The heat scale bar shows the Pearson correlation coefficient (*r*). The asterisks within the matrix show the significance threshold taken from the *P*-value for each comparison: *** = *P* < 0.001, ** = *P* < 0.01 and * = *P* < 0.05 created using R package corrplot (Wei and Simko, 2017). B) Ranked BLUEs for grain yield and C) grain width from a mixed linear model fitted via lme4 (Bates et al., 2015). Post-hoc pairwise comparisons of the BLUEs used the package emmeans (Lenth 2020). The asterisks indicate significant difference to ‘Paragon’ (*P* < 0.05). All error bars shown represent standard error of the predicted means. D) A photograph showing diversity in grain width using samples taken from plots in the NIAB_D field trial. The CSSLs with the three highest and lowest grain widths are shown, with ‘Paragon’ included for comparison.

### Introgression-trait association and validation

For the introgression-trait analysis, markers that were excluded during the earlier stages of QC were included to provide further resolution on introgression boundaries. The presence of awns was detected in a single NIAB_AB line (5A.07; **Figure 4A**). The CSSLs with neighbouring introgressions (5A.06 and 5A.08) were non-awned (**Figure 4A**). The awn inhibitor *B1* has been previously identified (TraesCS5A02G542800, 5A; 698,528,636 to 698,529,001bp; Huang *et al*., 2020). The introgression from TTD-140 in 5A.07 overlapped this gene (570 Mb to 700 Mb), whereas in 5A.06 and 5A.08 the introgressions were in adjacent regions (**Figure 4B**). The SNP marker AX-94504627 had a physical location (698,003,176 bp) close to the *B1* locus. The SNP cluster plot extracted from the Axiom Analysis Suite (Thermo Fisher Scientific) showed that the only CSSL to carry the TTD-140 allele at this location was 5A.07. As shown in **Figure 1**, NIAB_AB_5A.07 exhibits few off target introgressions on other chromosomes. No introgressions were observed in the regions harbouring the other reported awn inhibitors (Hooded locus (*Hd*)) on 4A and the awn inhibitor (*Tipped 2*) B2 on 6B (Mcintosh *et al*., 2013).

**Figure 4.**
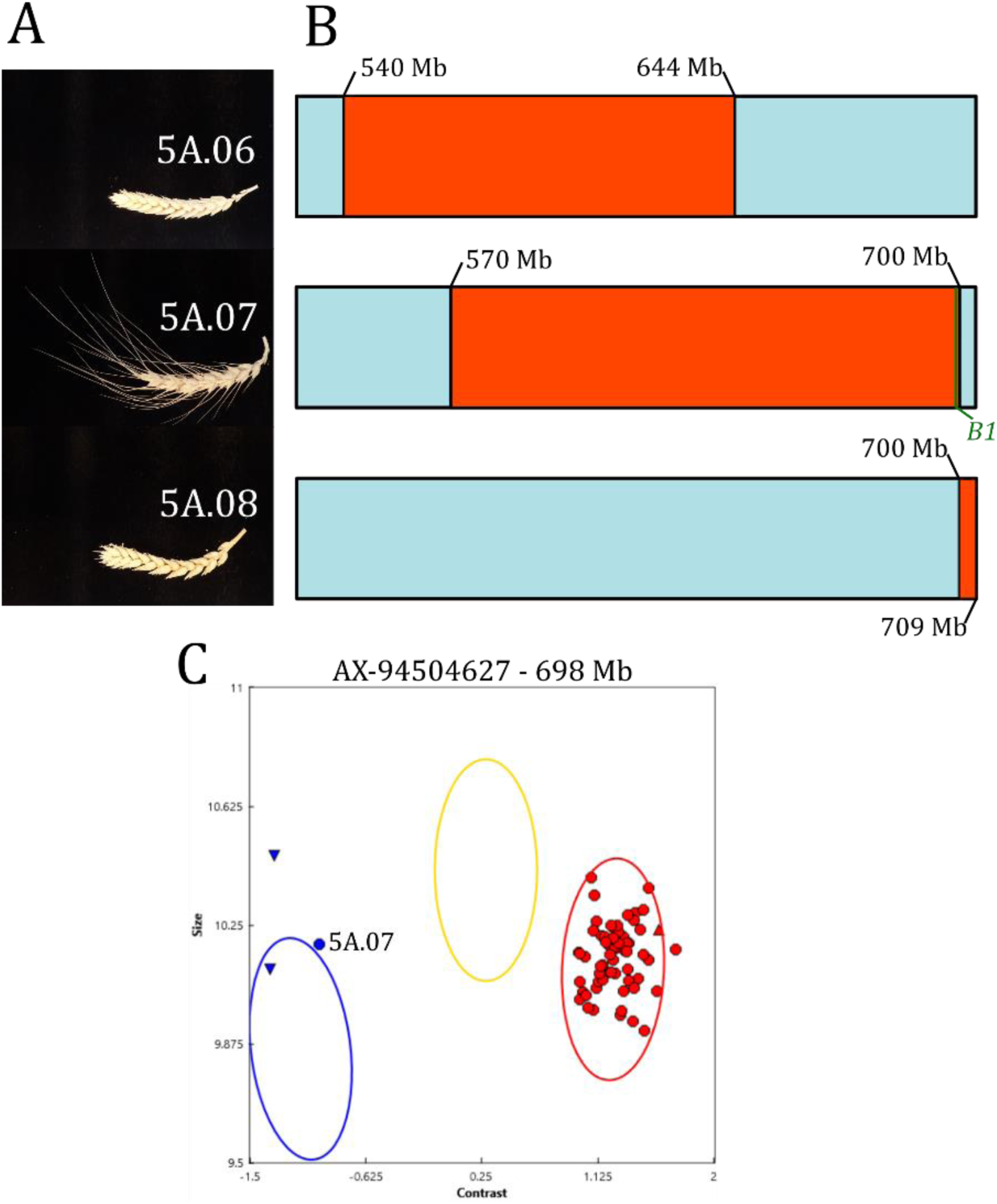
Introgression-trait association using the NIAB_AB population. A) Ears from the NIAB_AB glasshouse trial, showing the single awned CSSL (5A.07). B) The awned CSSL had an introgression from TTD-140 on 5A that extended over the *B1* awn inhibitor locus (Huang *et al.,* 2020) and awnless CSSLs 5A.06 and 5A.08 did not have an introgression in this region. A TTD-140 introgression is represented by the colour orange, light blue represents a ‘Paragon’ genotype. C) A genotyping plot taken from Axiom Analysis Suite (Thermo Fisher Scientific) for a SNP close to the gene (AX-94504627; 698,003,176 bp) showing CSSLs from the AB graphical genotype (circular points), two TTD-140 replicates (inverted triangle points) and ‘Paragon’ (a single triangle point). The only line to share the TTD-140 allele (blue) is CSSL.5A.07. Physical positions are in mega base pairs (Mb) and were taken from IWGSC (2018).

Three NIAB_D CSSLs displayed non-glaucous ear, stem and leaf phenotypes based on visual and RGB image assessment: 2D.01, 2D.02 and 3D.04. The CSSLs 2D.01 and 2D.02 were selected for NIAB-SHW041 introgressions at the start of 2D (**Figure 5A**), while 3D.04 was selected for a 3D introgression. However, 3D.04 also carried an off-target introgression at the start of 2D, from 1.6 to 13.7 Mb (**Figure 1**). The CSSL lines 2D.01 and 2D.02 carried an introgression on 2D from 2.7 to 17.6 Mb and 1.6 to 17.6 Mb, respectively. Additionally, 2D.02 had a downstream heterozygous introgression starting at around 19.7 Mb (**Figure 5A**). The

**Figure 5.**
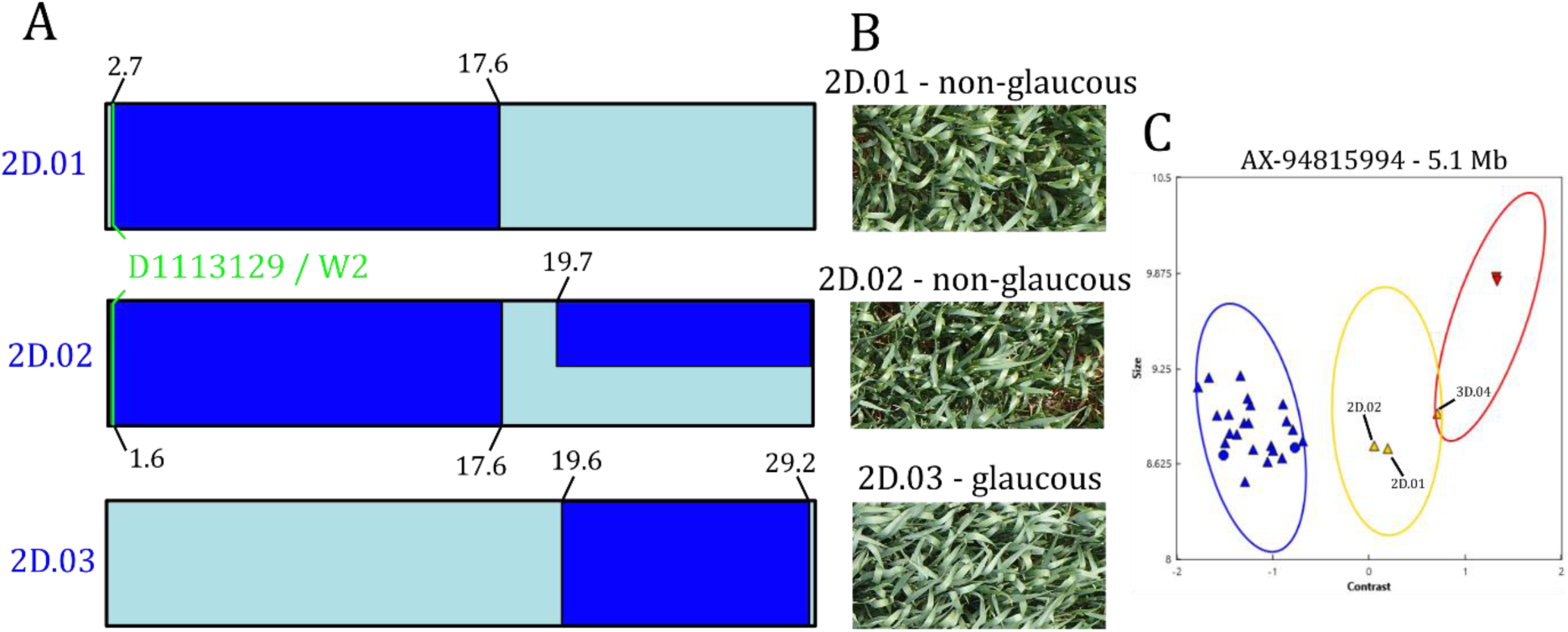
A) Schematic showing regions of chromosome 2D on three NIAB_D genome lines; light blue indicates a ‘Paragon’ genotype and dark blue indicates an introgression from NIAB-SHW041. The location of a putative QTL (linked to the marker D1113129) proposed to correspond to a known locus for wax production *Iw2* (Würschum *et al,.* 2020a) is overlayed on the plot. B). Canopy photos from the NIAB_D field trial show 2D.01 and 2D.02 were non-glaucous. C) A plot extracted from the genotyping software Axiom Analysis Suite (Thermo Fisher Scientific) for a SNP (AX-94815994; 5.1 Mb on 2D) that shows CSSLs genotypes from the D graphical genotype (triangle points), two of the D genome donor ENT-336 replicates (inverted triangle points) and ‘Paragon circular points). The three CSSLs that were non-glaucous in the NIAB_D field trial are labelled. Physical positions are shown in mega base pairs (Mb) taken from (IWGSC 2018).

CSSL 2D.03 did not share the non-glaucous phenotype, which is visible in the RGB canopy photographs of field plots for these genotypes (**Figure 5B**). The line 2D.03 had a homozygous introgression from 19.6 to 29.2 Mb. The previously identified putative marker-trait association with flag leaf glaucousness was based on a marker located at 2.9 Mb on 2D and hypothesized to be *Iw2* (Würschum *et al.,* 2020a), the D-genome paralog of *INHIBITOR of WAX1* (*Iw1*; identified by Huang *et al*., 2017). All three NIAB_D lines that were non-glaucous in the graphical genotype carried an introgression overlapping this region, which can be observed for 2D.01 and 2D.02 in **Figure 5A**. A SNP marker located at 5.1 Mb also confirmed that the three non-glaucous CSSLs did not possess the ‘Paragon’ allele close to the marker-trait association (**Figure 5C)**. The introgressions in these lines at the start of 2D were typically homozygous (**Figure 1**). However, the genotype calls at AX-94815994 are heterozygous (**Figure 5C)**. It is likely that the calls for the lines were homozygous for the ENT-336 (D-genome) allele but differences in ploidy impacting SNP calling between the CSSLs and the *Ae. tauschii* donor has separated the cluster. Whole genome graphical genotypes were inspected visually and the CSSLs 2D.01, 2D.02 and 3D.04 did not share any other overlapping introgressions.

## Discussion

We report the creation of a genetic resource for detecting and exploiting wheat progenitor variation. Publicly available pure seed, phenotyping and genotyping data support use as introgression donors and/or near-isogenic lines. The introgressions are genetically tractable, of reasonable size for forward use and have low off-target segments. This builds on early work to develop wheat resources for gene localisation and characterisation (Sears 1953 & 1954) and genetic mapping (Zemetra *et al*., 1986). Since this time numerous forward and reverse wheat genetic resources have been developed and the reported wheat CSSL series adds a further precision resource for exploiting progenitor diversity.

The NIAB_AB and NIAB_D series were created using backcrossing and MAS followed by high-density genotyping to facilitate progenitor segment introgression with low off-target effects. This approach is relatively time and cost intensive. Different methods of population construction have been proposed, including those with a lower marker requirement (e.g. Bian *et al*. (2010) used 8 informative markers at BC3F4 followed by pooled genotyping to extract individuals with small donor segments). This reduced genotyping approach is an alternative, low-cost approach to support CSSL development although larger numbers of individuals need to be progressed through backcrossing. Future development of CSSL series is likely to benefit from the reducing costs of genotyping, allowing more intensive screening and enhancing both accuracy of donor introgression and background recovery. The reducing cost of whole genome skim sequencing is also likely to allow higher resolution of introgression segments.

In both the NIAB_AB and NIAB_D series, we detected significant trait variation and transgressive segregation compared to ‘Paragon’. Grain size and shape are largely independent traits and we observed strong variation for grain length and width. Grain length of ancestral wheat has been shown to be generally higher than in elite bread wheat (Gegas *et al.,* 2010) and selection of alleles that confer increased grain length could increase grain size. Brinton *et al*. (2017) identified a significant QTL for TGW on chromosome 5A (*Qtgw-cb.5A*) demonstrating that the increase in grain area (driving TGW) was predominantly conferred by an increase in grain length. Yang *et al*. (2019) also identified a novel allele associated with increased grain length on 5AL (cloning *TaGL3-5A-G*). This allele, found in ancestral wheats, represents a breeding target, particularly if it could be pyramided in combination with favourable alleles for grain width (e.g. *TaGW2*; Simmonds *et al.,* 2016) and grain size (*TaGS5-3A*; Ma *et al.,* 2015). In the NIAB_AB population, there was a significant positive effect of grain yield per plant correlated with grain length. This identified individual CSSL lines with introgressions associated with increased grain length which are effectively near-isogenic lines for the TTD- 140 segment. The introgression in the lines with the greatest increase in grain length is in the same region as *TaGL3-5A* (Yang *et al.,* 2019) and future work could establish whether it contains the favourable allele described by Yang *et al* (2019). These lines could be field tested to validate the effect, and the segments transferred to additional elite backgrounds to test stability. In addition, the extreme positive and negative (compared to ‘Paragon’) lines could be used in a bulk-segregant genotyping approach (e.g. based on exome sequencing; Martinez *et al*., 2020 or RNA-seq; Zhu *et al.,* 2020) to determine the specific grain length controllers.

For the NIAB_D population, significant trait variation and high trait heritabilities were observed. The significant genotypic effects, despite the high recovery of ‘Paragon’ background, indicate effective capture (and contribution) of functional variation from the *Ae. tauschii* donor ENT-336. Variation was observed for grain yield although was only significant for CSSLs with yields lower than ‘Paragon’. Although not statistically significant, five CSSLs yielded higher than ‘Paragon’. Further testing through multi-site trials could establish if there is a significant yield effect associated with these CSSLs. Grain width was highly correlated with yield. This allowed identification of CSSLs with positive and negative effects for grain width which are available as near-isogenic lines, introgression segment donors and for further genetic characterisation, as above. Grain width has been well characterised in hexaploid wheat (Simmonds *et al.,* 2016) and alternative alleles for *TaGW2* can increase grain width and therefore grain weight, though phenotypes may be subtle due to allele dosage effects. Previous GWAS analysis of *Ae. tauschii* collections have identified numerous QTL for grain characteristics conferred by genetic effects on multiple chromosomes (Arora *et al*., 2017; Zhao *et al.,* 2021). The further development of markers for the regions underlying these QTLs will facilitate the validation of allelic effects and potential use for breeding.

In addition to identification of lines with positive and negative introgression effects, specific trait-introgression effects could be assessed based on physical locations of the segments. Although the series can’t be used directly for mapping *per se* due to the low power of detection it can be used to associate introgressions with phenotypes of interest. This is useful to validate previously observed effects as well as being a starting point for fine mapping. Here, our definition of introgression segment boundaries is limited to the coverage of Axiom® Wheat Breeder’s Genotyping Array (Allen *et al.,* 2017) but further genotyping would allow higher resolution and definition of boundaries and allow full automation of the process (array data requires manual curation below software quality thresholds). However, the current resolution is likely to be sufficient for transfer of segments via MAS.

The recurrent parent ‘Paragon’ carries the *B1* awn inhibition locus. Using the NIAB_AB population, and through replacement of the region with the wild type (recessive) progenitor allele via introgression we confirmed the location of the locus. This was near the edge of the introgression boundary but could be reliably detected and linked to the phenotype. The *Tipped 1* (*B1*) mutant is one of the three major loci linked to awn presence in wheat (Watkins and Ellerton, 1940; Sourdille *et al*., 2002; Yoshioka *et al.,* 2017) and has been identified as an important determinant of awn development (Mackay *et al*., 2014; Würschum *et al*., 2020b). The *B1* locus has been fine mapped in multiple studies to 5AL (DeWitt *et al*., 2020; Huang *et al*., 2020; Niu *et al*., 2020; Wang *et al*., 2020; Würschum *et al*., 2020b). Awn presence and the *B1* locus have been recently linked to an offset in the negative association between grain yield and protein content (Scott *et al*., 2021). The detection of the awn inhibitor segment demonstrates the use of the resource for categorical traits and could be used to further understand awn and linked-awn trait haplotype contributions from *T. diccocoides* and expedite their transfer to breeding. CSSLs with and without awns could also be used to further understand the mechanisms of the recently reported trade-offs between awns, quality, and nutritional traits (Scott *et al*., 2021).

In the NIAB_D population we demonstrate the use of trait-introgression association to validate a putative association with a categorical trait contributed by the progenitor *Ae. tauschii*. This used a step-by-step statistical approach enabling validation of a marker-trait association for glaucousness which fell slightly below the significance threshold in Würschum *et al*. (2020a). Our approach aligned the glaucous phenotype of the CSSL introgression segment with the physical position of the peak marker reported by Würschum *et al*. (2020a). In addition to confirming the genetic effect, this provides a 35K Axiom marker to facilitate MAS and tracking in breeding, adding to the previously identified DArTseq peak marker from Würschum *et al*. (2020a). This is likely to be advantageous given the relative ease of conversion of array probes compared to sequence-based probe conversion (Burridge *et al*., 2018).

Here we report the development, characterization, and initial testing of a CSSL wheat series as a tractable genetic resource adding to the existing portfolio of functional genomics resources for wheat improvement. The use of ‘Paragon’ as a genetic background complements other available wheat resources including the A.E. Watkins hexaploid wheat nested association mapping panel, and EMS and γ mutant populations (Wingen *et al*., 2017), QTL datasets (Przewieslik-Allen *et al*., 2021), and publicly available synthetic hexaploid wheat populations (Adamski *et al*., 2020). Further work is required to reduce off-target introgressions and to split multiple introgression lines into discreet single-introgression lines. The application of higher density markers and/or sequencing will support further precision in defining introgression boundaries. The extreme phenotypic lines can be further validated as near isogenic lines and used as donor lines for transfer of introgressions to additional elite parents for use in breeding. The NIAB_AB and NIAB_D lines can also be intercrossed to create favourable introgression complements (e.g. grain length and grain width) for further testing and development of targeted multi-introgression lines for specific traits of interest. Overall, the resource reported here increases access to wheat genomic resources for research and breeding improvements.

## Acknowledgements

Development of the CSSL resource was supported by the UK’s Biotechnology and Biological Sciences Research Council via the Wheat Improvement Strategic Program (BB/I002561/1) and Designing Future Wheat (BB/P016855/1). We thank Professor Andy Greenland, Dr Nicolas Gosman and Dr Keith Gardner for their contribution and insights into the development of the resource. We thank Michael Scott for supplying the BLAST+ output for the SNP markers and Dr Rob Davey for facilitating trial and image uploads to Grassroots.

## Author contributions

P.H., C.U., K.J.E. and A.R.B. conceived the project; R.H., F.J.L, T.I.C.W., A.R.B designed experiments; R.H., F.J.L, A.J.B., A.L., A.M.P.-A. performed the experiments; F.J.L, T.I.C.W., P.H. and A.R.B analysed experiments, R.H., F.J.L, T.I.C.W. and A.R.B wrote the manuscript with inputs from all other authors.

## Data availability

Seed of the germplasm reported here is directly available from the Germplasm Resources Unit at the John Innes Centre (via https://www.seedstor.ac.uk/), including the parental line ‘Paragon’ (WCSSL0001), donor lines TTD-140 (WCSSL0002) and NIAB-SHW041 (WCSSL0003) and for the NIAB_AB (WCSSL0004 to WCSSL0080) and NIAB_D (WCSSL0081 to WCSSL0112) populations.

The Axiom® Wheat Breeder’s Genotyping Array data is available for direct download for each of the three genomes here:

https://www.cerealsdb.uk.net/cerealgenomics/CerealsDB/genotyping_data/NIAB_CSSL_Agenome.csv https://www.cerealsdb.uk.net/cerealgenomics/CerealsDB/genotyping_data/NIAB_CSSL_Bgenome.csv https://www.cerealsdb.uk.net/cerealgenomics/CerealsDB/genotyping_data/NIAB_CSSL_Dgenome.csv

Plant images from the NIAB_AB glasshouse trial are available from: https://opendata.earlham.ac.uk/wheat/under_license/toronto/Horsnell,Leigh,Bentley,Wright_2020_NIAB_CSSL_D_genome_yield_trial_H2020/

Plot images from the NIAB_D field trial are available from: https://opendata.earlham.ac.uk/wheat/under_license/toronto/Horsnell,Leigh,Bentley,Wright_2020_NIAB_CSSL_D_genome_yield_trial_H2020/

Data from the NIAB_D field trial are available from: https://grassroots.tools/fieldtrial/study/61faaf25c68884365e7bcc34

