## Supplementary Figures for "A wheat chromosome segment substitution line series supports characterisation and use of progenitor genetic variation"

**Supplementary Figure S1.** Schematic of crossing scheme used to develop the two complementary CSSL series: NIAB_AB and NIAB_D populations.

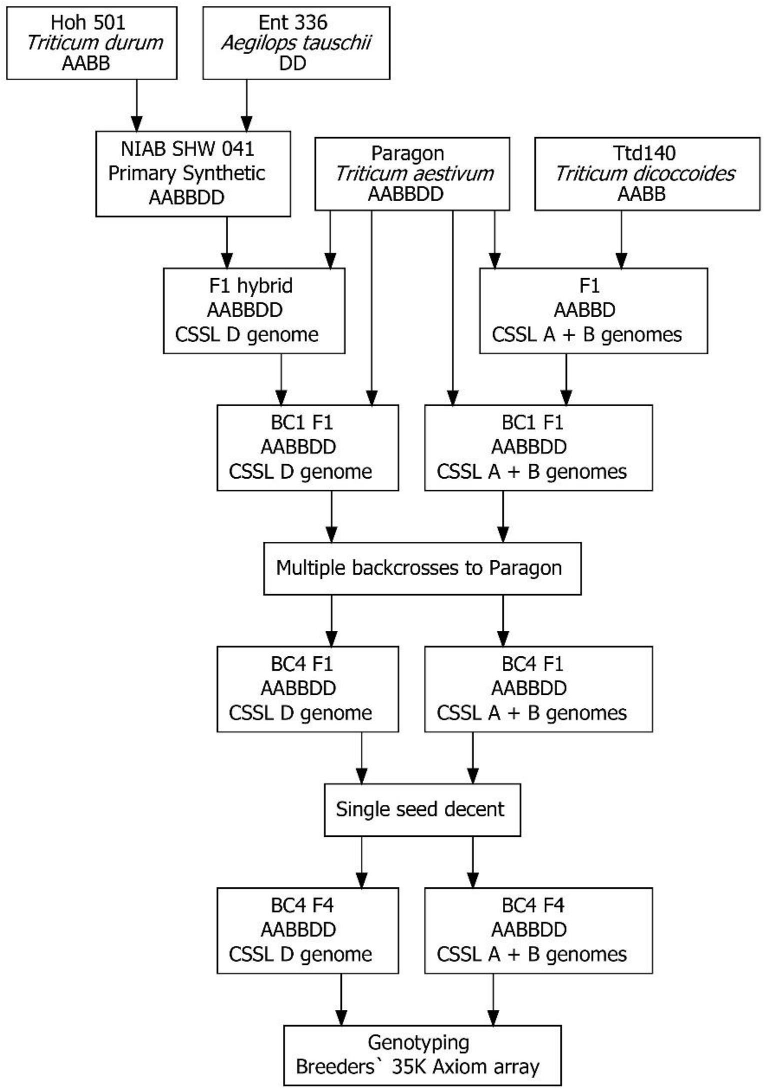

**Supplementary Figure S2.** Principle coordinate analysis (PCoA) characterises the genetic diversity between parents of the two series using the first two axis from each PCoA with the percentage variation captured by the eigenvalues shown. A) The relationship between 'Paragon' and a collection of different tetraploid wheat species. The plot includes T. dicoccoides introgression donor TTD-140 and uses markers mapped to the A and B genomes. B) The relationship between 51 primary NIAB synthetic hexaploid wheats and 'Paragon', this includes the introgression donor line NIAB-SHW041. Only D genome markers were used in the comparison. Marker genome assignments are from the consensus genetic linkage map from Allen et al. (2017). The PCoA was completed on Euclidean genetic distance matrices and implemented using the R package ape (Paradis & Schliep 2018).

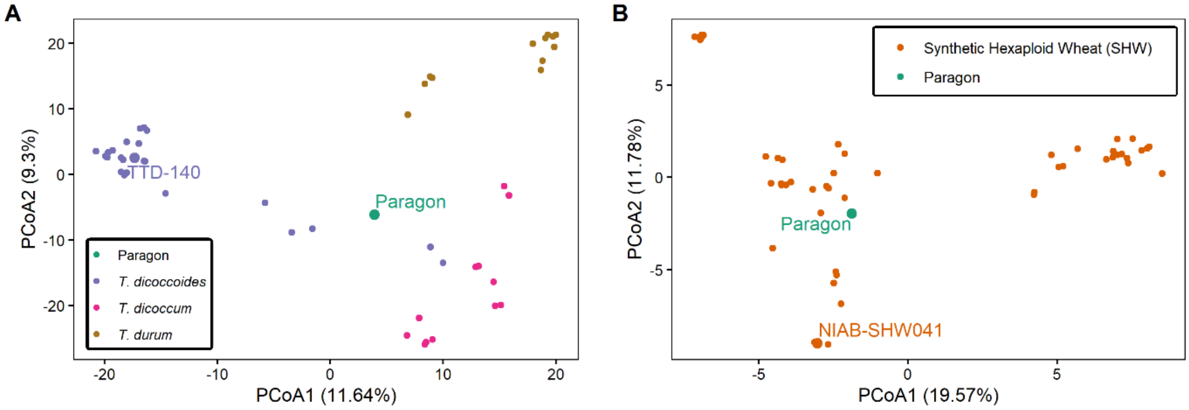

**Supplementary Figure S3.** Complete graphical genotypes for the NIAB_AB CSSLs selected for TTD-140 introgressions on the A genome created using the R package SelectionTools (v-19.1, www.uni-giessen.de). This displays graphical genotypes for the targeted A genome and the off-target B and D genomes. Chromosomes were scaled on genetic distance using 4,990 SNPs and genetic position from the 35K array consensus map (Allen et al., 2017). Light blue represents a marker call with a 'Paragon' allele and dark red represents the alternative marker allele. White spaces show missing data and heterozygous markers are shown by a split of the colours.

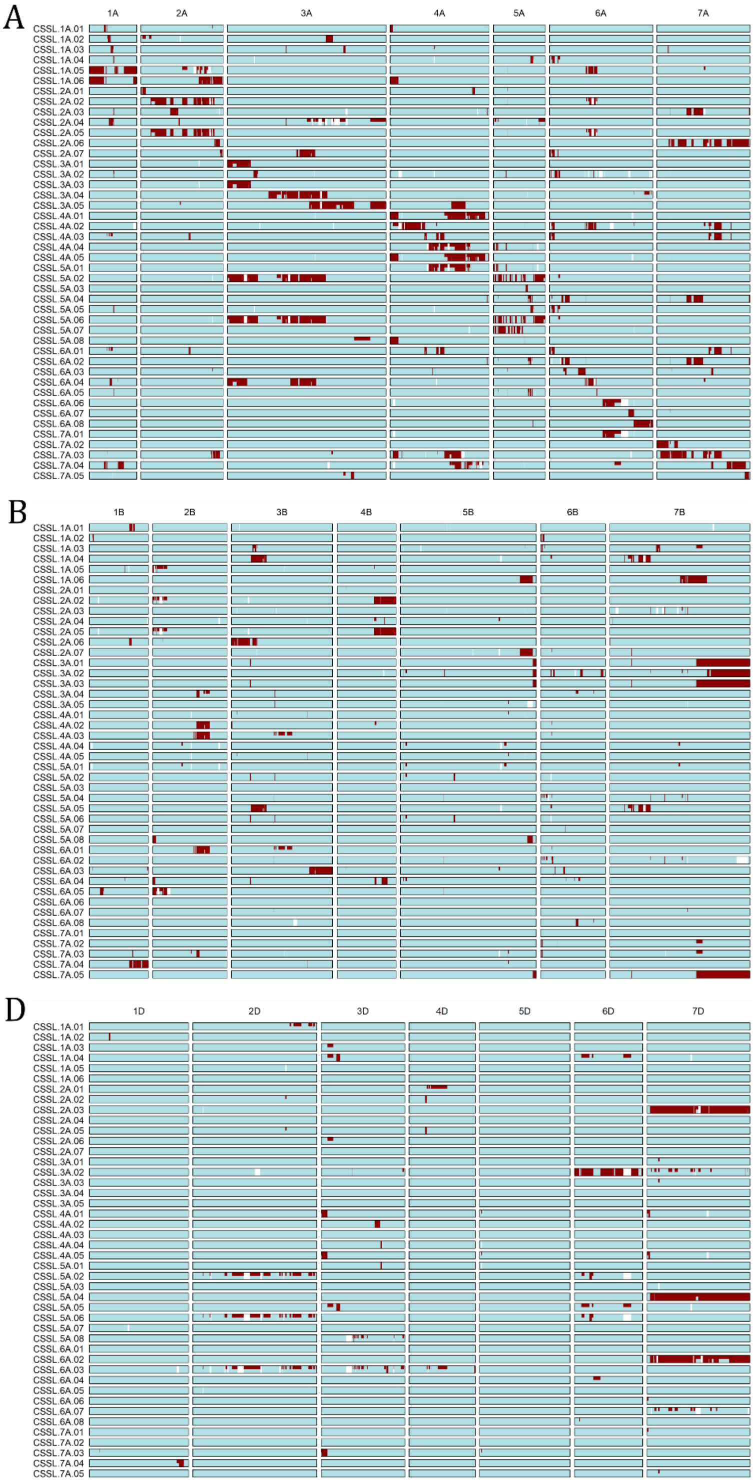

**Supplementary Figure S4.** Complete graphical genotypes for the NIAB_AB CSSLs selected for TTD-140 introgressions on the B genome created using the R package SelectionTools (v-19.1, www.uni-giessen.de). This displays graphical genotypes for the targeted B genome and the off-target A and D genomes. Chromosomes are scaled on genetic distance using 4,983 SNPs and genetic positions from the 35K array consensus map (Allen et al., 2017). Light blue represents a marker call with a 'Paragon' allele and dark red represents the alternative marker allele. White spaces show missing data and heterozygous markers are shown by a split of the colours.

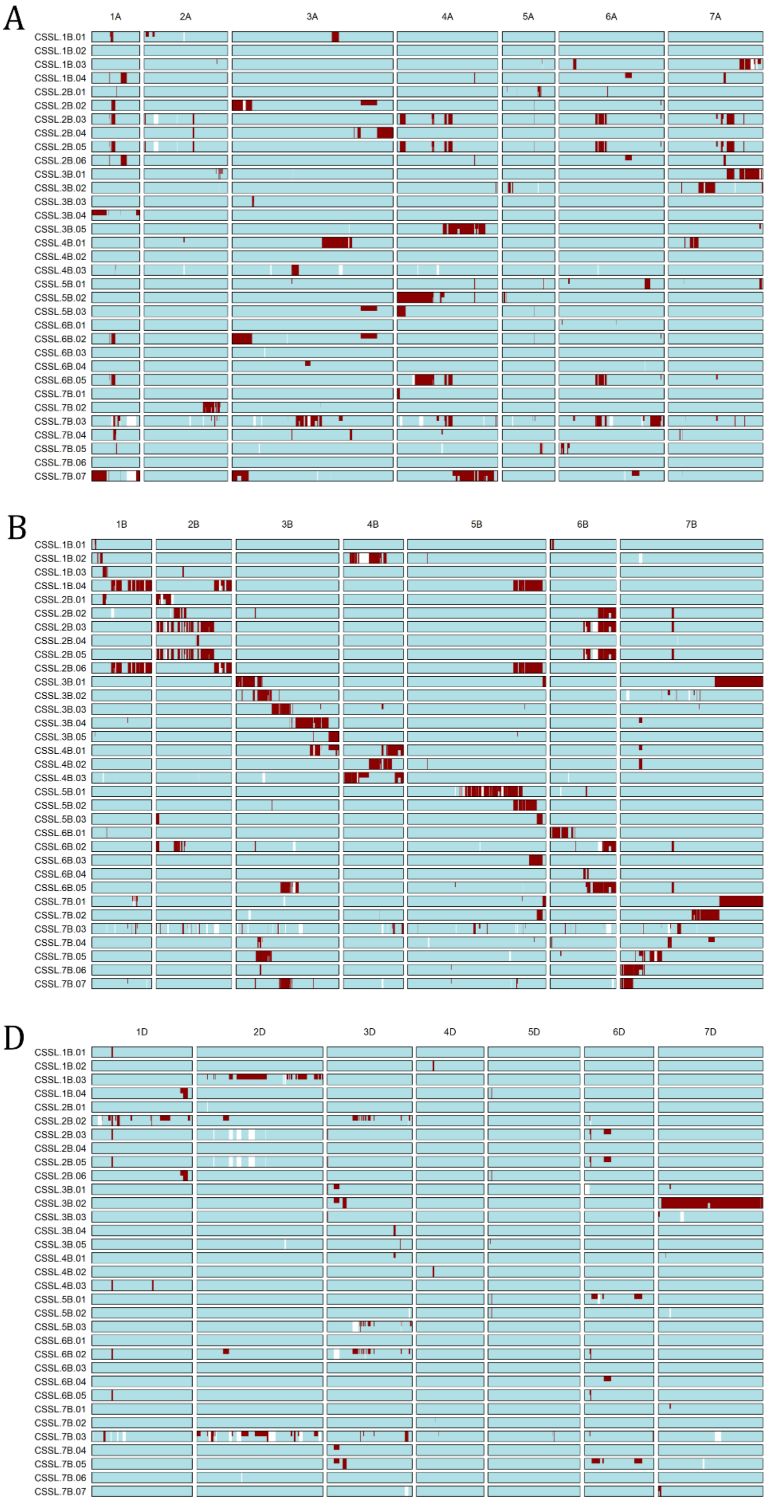

**Supplementary Figure S5.** Complete graphical genotypes for the NIAB_D CSSLs selected for NIAB-SHW041 introgressions on the D genome created using the R package SelectionTools (v-19.1, www.uni-giessen.de). This displays graphical genotypes for the targeted D genome and the off-target A and B genomes. Chromosomes are scaled on genetic distance using 4,982 SNPs and genetic position from the 35K array consensus map (Allen et al., 2017). Light blue represents a marker call with a ‘Paragon’ allele and dark red represents the alternative marker allele. White spaces show missing data and heterozygous markers are shown by a split of the colours.

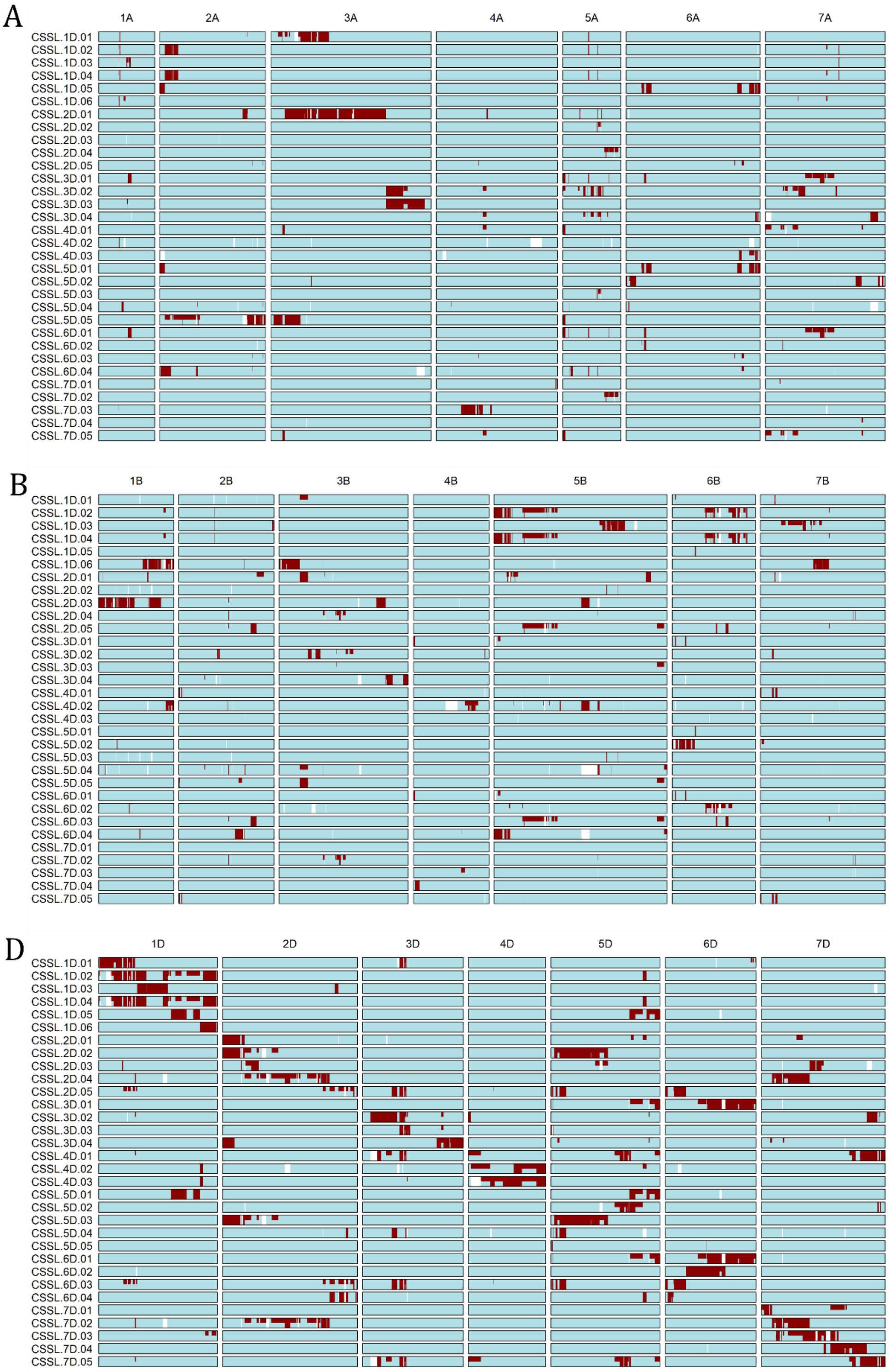
