## Supplementary Tables for "A wheat chromosome segment substitution line series supports characterisation and use of progenitor genetic variation"

**Supplementary Table S1.** The creation of the NIAB_AB and NIAB_D populations to BC4 generation involved creation of plants with genotyping at each stage to identify individuals to progress via marker-assisted selection.

| | Plants created | | Individuals genotyped | | Markers assayed | | Individuals selected | | Genotyping |
|---|---|---|---|---|---|---|---|---|---|
| Generation | AB | D | AB | D | AB | D | AB | D | |
| $F_1$ | 6 | 14 | | | | | | | |
| $BC_1F_1$ | 65 | 317 | 65 | 181 | 142 | 150 | 13 | 14 | KASP |
| $BC_2F_1$ | 155 | 553 | 155 | 186 | 189 | 150 | 35 | 34 | KASP |
| $BC_3F_1$ | 690 | 753 | 298 | 278 | 201 | 150 | 83 | 44 | KASP |
| $BC_4F_1$ | 429 | 1134 | 354 | 338 | 159 | 148 | 138 | 51 | KASP |
| $BC_4F_2$ | 1041 | n/a | 1041 | 366 | 145 | 148 | 199 | 57 | KASP |
| $BC_4F_3$ | 1344 | n/a | 1344 | 54 | 124 | 148 | 152 | 33 | KASP |
| $BC_4F_4$ | 152 | n/a | 152 | 362 | 35K* | 35K* | 152 | 33 | 35K array |

*Genotyped using the Axiom® Wheat Breeder's Genotyping Array (Allen et al., 2017)

**Supplementary Table S2.** Selection of 98 A-genome and 93 B-genome specific KASP to use in marker-assisted backcrossing. Markers were selected to be evenly distributed and co-dominant (where possible) and polymorphic between the recurrent parent Paragon and the AB genome donor TTD-140 based on genetic mapping information from the Avalon x Cadenza population. All data available from https://www.cerealsdb.uk.net.

| | | | **SNP call** | |
|---|---|---|---|---|
| **Marker** | **Chromosome** | **cM** | **Paragon** | **TTD-140** |
| BS00022677 | 1A | 5.92 | G:G | A:A |
| BS00023201 | 1A | 16.2 | A:A | G:G |
| BS00078982 | 1A | 47.65 | A:A | C:C |
| BS00022275 | 1A | 60.1 | C:C | T:T |
| BS00023462 | 1A | 63 | G:G | A:A |
| BS00022207 | 1A | 64.2 | T:T | G:G |
| BS00077990 | 1A | 75.6 | C:C | T:T |
| BS00023203 | 1A | 105.5 | T:T | C:C |
| BS00053661 | 2A | 30.2 | A:A | C:C |
| BS00079036 | 2A | 38.67 | T:C | C:C |
| BS99999945 | 2A | 51.6 | A:A | G:G |
| BS00075524 | 2A | 51.6 | T:T | C:C |
| BS00045900 | 2A | 58.8 | A:A | G:G |
| BS00059607 | 2A | 80.9 | C:C | G:G |
| BS00078489 | 2A | 83.4 | T:T | C:C |
| BS00035883 | 2A | 90.8 | T:T | G:G |
| BS00082084_51 | 2A | 105.5 | C:C | T:T |
| BS00022381 | 2A | 105.5 | C:C | T:T |

| | | | | |
|---|---|---|---|---|
| BS99999939 | 2A | 126.0 | C:C | T:T |
| BS00081630 | 2A | 126 | A:A | C:C |
| BS00063368 | 2A | 133 | T:T | C:C |
| BS00107316 | 2A | 146.7 | G:G | T:T |
| BS00007689 | 2A | 160.2 | T:T | C:C |
| BS00069105 | 2A | 171.5 | T:T | C:C |
| BS00084881 | 2A | 186.1 | A:A | G:G |
| BS00022985 | 3A | 0 | C:C | G:G |
| BS00073748 | 3A | 1 | A:A | G:G |
| BS00013584_51 | 3A | 57.2 | A:A | G:G |
| BS00025191 | 3A | 61.7 | T:T | G:G |
| BS00032524 | 3A | 62 | T:T | C:C |
| BS00037537 | 3A | 74.24 | G:G | A:A |
| BS00070014 | 3A | 83.7 | A:A | G:G |
| BS00077756 | 3A | 88.59 | T:T | C:C |
| BS00022658 | 3A | 96.89 | G:G | A:A |
| BS00024548 | 3A | 117.16 | G:G | A:A |
| BS00022968 | 3A | 138.3 | C:C | T:T |
| BS00022735 | 3A | 163.78 | G:G | A:A |
| BS00043286 | 4A | 1.3 | A:A | G:G |
| BS00065863 | 4A | 6.9 | G:G | T:T |
| BS00065607 | 4A | 13.5 | A:A | G:G |
| BS00022169 | 4A | 28.7 | A:A | G:G |
| BS00049911 | 4A | 29.23 | G:G | T:T |
| BS00088726 | 4A | 64.97 | A:A | G:G |
| BS00021989 | 4A | 112.49 | G:G | A:A |
| BS00099534 | 5A | 3.2 | A:A | G:G |
| BS00034304 | 5A | 12.71 | C:C | T:T |
| BS00021939 | 5A | 17.6 | G:G | A:A |
| BS00077990 | 5A | 23 | G:G | A:A |
| BS00041220 | 5A | 30.8 | T:T | C:C |
| BS00023532 | 5A | 41.35 | T:T | G:G |
| BS00066143 | 5A | 50.4 | T:T | C:C |
| BS00022683 | 5A | 59.8 | T:T | C:C |
| BS99999943 | 5A | 79.9 | A:A | G:G |
| BS00077855 | 5A | 81.5 | C:C | T:T |
| BS00098172 | 5A | 91.1 | C:C | T:T |
| BS00072754 | 5A | 105.1 | A:A | T:T |
| BS00042336 | 5A | 122.5 | G:G | A:A |
| BS00063793_51 | 5A | 130.0 | A:A | G:G |
| BS00022891 | 5A | 159.6 | C:C | A:A |
| BS00022028 | 5A | 168 | T:T | A:A |
| BS00029462 | 5A | 169.4 | T:T | C:C |
| BS00021969 | 5A | 192.4 | T:T | C:C |
| BS00022664 | 5A | 197.79 | C:C | G:G |
| BS00022517 | 6A | 1 | T:T | C:C |

| | | | | |
|---|---|---|---|---|
| BS00022584 | 6A | 1 | A:A | G:G |
| BS00094998 | 6A | 4.9 | C:C | G:G |
| BS00083630 | 6A | 12.5 | T:T | G:G |
| BS99999934 | 6A | 49.5 | G:G | A:A |
| BS00085980 | 6A | 53.38 | A:A | G:A |
| BS00093964 | 6A | 59.96 | G:G | A:A |
| BS99999944 | 6A | 75.1 | G:G | A:A |
| BS00065852 | 6A | 75.2 | T:T | C:C |
| BS00031178 | 6A | 89.94 | C:C | T:T |
| BS00033795 | 6A | 98.8 | C:C | T:T |
| BS00022628 | 6A | 105.96 | T:T | C:C |
| BS00021961 | 6A | 115.74 | G:G | A:A |
| BS00029954 | 6A | 115.86 | A:A | T:T |
| BS00074754 | 6A | 122 | T:T | C:C |
| BS00021965 | 6A | 137.91 | G:G | A:A |
| BS00023088 | 6A | 146 | T:T | A:A |
| BS99999946 | 7A | 1.8 | C:C | T:T |
| BS00022386 | 7A | 12.58 | G:G | A:A |
| BS00012880 | 7A | 23.9 | T:T | C:C |
| BS99999940 | 7A | 28.0 | C:C | A:A |
| BS00022696 | 7A | 52 | A:A | G:G |
| BS00022097 | 7A | 78.57 | A:A | G:G |
| BS00023225 | 7A | 110 | A:A | C:C |
| BS99999936 | 7A | 120.9 | T:T | C:C |
| BS00068589 | 7A | 137.7 | T:T | C:C |
| BS00030391 | 7A | 148 | T:T | C:C |
| BS00031028 | 7A | 148.26 | A:A | G:G |
| BS00071478 | 7A | 150.1 | T:T | C:C |
| BS00002510 | 7A | 165.5 | C:C | T:T |
| BS00023218 | 7A | 176.1 | G:G | C:C |
| BS00033613 | 7A | 191.15 | C:C | T:T |
| BS00110699 | 7A | 217.26 | G:G | C:C |
| BS00022137 | 7A | 229.1 | A:A | G:G |
| BS00023002 | 7A | 332 | T:T | G:G |
| BS00022564 | 1B | 1 | A:A | G:G |
| BS00070139 | 1B | 5.8 | A:A | C:C |
| BS00022571 | 1B | 18.4 | G:G | C:C |
| BS00082143 | 1B | 29.3 | G:G | C:C |
| BS00021941 | 1B | 45.1 | A:A | C:C |
| BS00022539 | 1B | 45.11 | T:T | C:C |
| BS00022609 | 1B | 56.67 | T:T | G:G |
| BS00066165 | 1B | 59.8 | G:G | A:A |
| BS00022822 | 1B | 60 | C:C | T:T |
| BS00022093 | 1B | 87.94 | G:G | C:C |
| BS00041355 | 1B | 91.08 | C:C | T:T |
| BS00022851 | 1B | 97.16 | A:A | G:G |

| | | | | |
|---|---|---|---|---|
| BS00069615 | 1B | 100.9 | T:T | G:G |
| BS00077232 | 1B | 107 | G:G | C:C |
| BS00035268 | 1B | 135.89 | C:C | T:T |
| BS00078414 | 1B | 151.3 | G:G | T:T |
| BS00060686 | 1B | 174 | G:G | T:T |
| BS00084668 | 2B | 10.8 | A:A | G:G |
| BS00070900 | 2B | 20.1 | G:G | A:A |
| BS00022950 | 2B | 43.6 | G:G | A:A |
| BS00038820 | 2B | 61.1 | T:C | C:C |
| BS00023097 | 2B | 86.2 | G:G | C:C |
| BS00041922 | 2B | 91.42 | T:T | C:C |
| BS00047070 | 2B | 95.5 | A:A | G:G |
| BS00022374 | 2B | 102.1 | T:T | C:C |
| BS00022940 | 2B | 114 | A:A | G:G |
| BS00013126 | 2B | 118.84 | T:T | A:A |
| BS00071171 | 2B | 140.8 | C:C | G:G |
| BS00064776_51 | 3B | 0.0 | T:T | C:C |
| BS00024783 | 3B | 2.4 | A:A | G:G |
| BS00026471 | 3B | 29.72 | G:G | T:T |
| BS00064778 | 3B | 31.12 | G:G | A:A |
| BS99999931 | 3B | 44.1 | A:A | G:G |
| BS00070456 | 3B | 55.93 | G:G | A:A |
| BS00022961 | 3B | 77 | T:T | C:C |
| BS00062676 | 3B | 78 | G:G | A:A |
| BS00079014 | 3B | 90.3 | G:G | A:A |
| BS00022316 | 3B | 105.44 | G:G | A:A |
| BS00022403 | 3B | 114.52 | A:A | G:G |
| BS00056737 | 3B | 118.9 | A:A | G:G |
| BS00022415 | 3B | 135.3 | G:G | T:T |
| BS00030651 | 3B | 139 | T:T | C:C |
| BS00071182 | 3B | 160 | T:T | C:C |
| BS00022441 | 3B | 172.3 | A:A | G:G |
| BS00062859 | 3B | 191.21 | C:C | G:G |
| BS00058946 | 4B | 11.4 | C:C | G:G |
| BS00044374 | 4B | 36.9 | A:A | C:C |
| BS00021984 | 4B | 47.5 | T:T | C:C |
| BS00024846 | 4B | 51.54 | T:T | C:C |
| BS00022194 | 4B | 60.7 | T:T | C:C |
| BS00076033 | 4B | 66.3 | A:A | G:G |
| BS00037380 | 4B | 68.95 | T:T | C:C |
| BS00091035_51 | 4B | 75.4 | G:G | A:A |
| BS00072221 | 4B | 85.5 | A:A | T:T |
| BS00022793 | 4B | 108.4 | A:A | G:G |
| BS00023204 | 4B | 110.16 | T:T | C:C |
| BS00023161 | 5B | 1 | C:C | T:T |
| BS00023163 | 5B | 2.4 | T:T | C:C |

| | | | | |
|---|---|---|---|---|
| BS00022555 | 5B | 4.6 | T:T | C:C |
| BS00022336 | 5B | 20 | T:T | C:C |
| BS00022231 | 5B | 32.1 | G:G | G:A |
| BS00106043 | 5B | 46.79 | T:T | C:C |
| BS00023127 | 5B | 51 | G:G | A:A |
| BS00108019 | 5B | 62.8 | T:T | C:C |
| BS00099385 | 5B | 64.59 | C:C | T:T |
| BS00023803 | 5B | 74 | G:G | A:A |
| BS00077690 | 5B | 79.6 | A:A | G:G |
| BS00022151 | 5B | 83.3 | A:A | T:T |
| BS00108020_51 | 5B | 88.5 | C:C | T:T |
| BS00065697 | 5B | 107.8 | G:G | A:A |
| BS00022400 | 5B | 123.4 | A:A | G:G |
| BS00022065 | 5B | 146.7 | A:A | G:G |
| BS00098520 | 5B | 169.02 | A:A | G:G |
| BS00022755 | 5B | 177.62 | A:A | G:G |
| BS00021948 | 5B | 186 | G:G | A:A |
| BS99999935 | 5B | 187.8 | A:A | G:G |
| BS00110474 | 5B | 205.3 | C:C | T:T |
| BS00093063 | 6B | 0.6 | C:C | T:T |
| BS00023196 | 6B | 30.9 | G:G | T:T |
| BS00107725 | 6B | 47.9 | G:G | C:C |
| BS00068615 | 6B | 61.3 | A:A | G:G |
| BS00022832 | 6B | 75.9 | T:T | G:G |
| BS00072715 | 6B | 81.48 | T:T | G:G |
| BS00093914 | 6B | 85.9 | C:C | G:G |
| BS00085589 | 6B | 101.24 | G:G | A:A |
| BS00076101 | 6B | 120.6 | T:T | G:G |
| BS00110651 | 6B | 127.8 | C:C | T:T |
| BS99999938 | 7B | 5.0 | G:G | A:A |
| BS00013057 | 7B | 8.74 | G:G | C:C |
| BS00022810 | 7B | 9.3 | A:A | G:G |
| BS00035559 | 7B | 31.1 | A:A | C:C |
| BS00081841 | 7B | 51.2 | A:A | G:G |
| BS00022175 | 7B | 71.01 | A:A | G:G |

**Supplementary Table S3.** Selection of 65 D-genome KASP to for background selection in NIAB_AB for individuals with D genomes from the hexaploid recurrent parent. Markers were selected to be evenly distributed and co-dominant (where possible) and specific to the D genome of the recurrent parent Paragon. Chromosome allocation and genetic distance (cM) were based on the Avalon x Cadenza (AxC) and Spark x Rialto (SxR) populations. All data available from https://www.cerealsdb.uk.net.

| SNP name | AxC_Chr | AxC cM | SxR_Chr | SxR cM | Paragon | TTD-140 |
|---|---|---|---|---|---|---|
| BS00055737 | 1D | 23.2 | 1D | 51.44 | T:T | - |

| | | | | | | |
|---|---|---|---|---|---|---|
| BS00103876 | 1D | 37.9 | | | G:G | - |
| BS00079462 | 1D | 47.3 | | | G:G | G:G |
| BS00022633 | 1D | 50.0 | | | C:C | - |
| BS00074825 | 1D | 50.6 | 1D | 74.18 | A:A | - |
| BS00063145 | 1D | 50.6 | 1D | 74.18 | C:C | - |
| BS00065478 | 1D | 50.6 | 1D | 74.18 | G:G | - |
| BS00022027 | 1D | 52.5 | | | C:C | - |
| BS00067768 | 1D | 57.8 | 1D | 73.1 | A:A | - |
| BS00082503 | 1D | 61.7 | 1D | 69.87 | G:G | - |
| BS00075001 | 1D | 62.8 | 1D | 69.87 | G:G | - |
| BS00040568 | 1D | 66.8 | | | T:T | - |
| BS00022485 | 1D | 133.5 | | | G:G | - |
| BS00099705 | 1D | 134.9 | | | C:C | - |
| BS00059043 | 2D | 14.3 | | | G:G | - |
| BS00062567 | 2D | 14.3 | | | T:T | - |
| BS00022976 | 2D | 57.5 | | | G:A | - |
| BS00021950 | 2D | 69.3 | | | G:G | - |
| BS00020492 | 2D | 73.0 | | | A:A | - |
| BS00023211 | 2D | 82.0 | 2D | 8.76 | G:G | - |
| BS00081578 | 2D | 87.2 | | | G:G | T:T |
| BS00064538 | | | 2D | 35.9 | C:C | - |
| BS00022942 | | | 2D | 170.26 | T:C | - |
| BS00023062 | | | 2D | 72.91 | T:T | - |
| BS00022305 | | | 2D | 72.91 | T:T | T:C |
| BS00049908 | | | 3D | 11.56 | T:T | - |
| BS00034812 | 3D | 2.05 | | | | |
| BS00037276 | 3D | 61.2 | | | G:G | - |
| BS00098495 | 3D | 73.2 | | | T:C | - |
| BS00031366 | 4D | 12.9 | | | G:G | - |
| BS00023046 | 4D | 16.6 | 4D | 16.12 | T:T | C:C |
| BS00023014 | 4D | 54.1 | 4D | 56.67 | A:A | - |
| BS00023131 | | | 4D | 16.12 | | |
| BS00023103 | | | 5D | 69.65 | T:T | - |
| BS00067650 | 5D | 125.3 | | | T:T | - |
| BS00088591 | 5D | 148.6 | | | T:T | - |
| BS00075107 | 5D | 149.1 | | | G:G | - |
| BS00022291 | 5D | 150.3 | | | - | A:A |
| BS00022036 | 5D | 150.3 | | | G:G | A:A |
| BS00022850 | 5D | 150.3 | | | G:G | - |
| BS00022157 | 5D | 151.7 | | | G:A | - |
| BS00058709 | 5D | 161.2 | 5D | 169.14 | T:T | - |
| BS00021991 | 5D | 161.8 | 5D | 169.14 | A:A | G:A |
| BS00022688 | 5D | 170.0 | 5D | 152.82 | C:C | - |
| BS00074353 | 5D | 170.0 | | | T:T | C:C |
| BS00085929 | 6D | 0.0 | | | A:A | G:G |
| BS00093153 | 6D | 54.1 | | | G:G | - |

| | | | | | | |
|---|---|---|---|---|---|---|
| BS00022856 | | | 6D | 61.21 | A:A | A:A |
| BS00080047 | 6D | 116.6 | 6D | 136.76 | C:C | - |
| BS00022204 | 6D | 116.6 | 6D | 136.76 | T:T | - |
| BS00022787 | | | 6D | 141.29 | G:G | - |
| BS00110152 | | | 7D | 64.05 | C:C | - |
| BS00110300 | | | 7D | 64.05 | C:C | - |
| BS00021987 | 7D | 27.7 | 7D | 122.95 | A:A | - |
| BS00023045 | 7D | 27.7 | 7D | 122.95 | A:A | - |
| BS00022721 | 7D | 27.7 | | | C:C | - |
| BS00022511 | 7D | 27.7 | 7D | 122.95 | T:T | T:T |
| BS00075780 | 7D | 28.3 | | | G:G | - |
| BS00096478 | 7D | 28.3 | 7D | 122.95 | G:G | - |
| BS00075898 | 7D2 | 1.1 | | | T:T | - |
| BS00062860 | 7D2 | 4.5 | | | G:G | - |
| BS00023150 | 7D3 | 46.2 | | | C:C | - |
| BS00070188 | 7D3 | 48.9 | | | G:G | - |
| BS00094533 | 7D3 | 48.9 | | | T:T | - |

**Supplementary Table S4.** Selection of 60 D-genome KASP markers for use in marker-assisted backcrossing. Markers were selected to be evenly distributed and co-dominant (where possible) and polymorphic between the recurrent parent Paragon and the D-genome *Ae. tauschii* donor ENT-336. Chromosome allocation and genetic distance (cM) were based on the Avalon x Cadenza (AxC) and Spark x Rialto (SxR) populations. All data available from https://www.cerealsdb.uk.net.

| | | | | | SNP calls | |
|---|---|---|---|---|---|---|
| **SNP ID** | **AxC_Chr** | **AxC cM** | **SxR_Chr** | **SxR cM** | **Paragon** | **ENT-336** |
| BS00022323 | 1D | 1.1 | | | A:A | C:C |
| BS00055737 | 1D | 23.7 | 1D | 50.1 | T:T | C:C |
| BS00118867 | 1D | 27.5 | | | C:C | T:T |
| BS00122008 | 1D | 35.8 | | | G:G | A:A |
| BS00078897 | 1D | 43.6 | | | T:T | C:C |
| BS00011451 | 1D | 46.8 | 1D | 73.7 | G:G | A:A |
| BS00082503 | 1D | 56.7 | 1D | 69.3 | G:G | A:A |
| BS00119899 | 1D | 56.7 | | | T:T | C:C |
| BS00040568 | 1D | 61.7 | | | T:T | C:C |
| BS00022188 | 1D | 79.2 | | | C:C | T:T |
| BS00089269 | 1D | 107.7 | | | C:C | T:T |
| BS00108448 | 2D | 5.1 | | | G:G | A:A |
| BS00059043 | 2D | 14.8 | | | G:G | C:C |
| BS00043986 | 2D | 30.0 | | | G:G | A:A |
| BS00093757 | 2D | 36.3 | | | T:T | C:C |
| BS00022976 | 2D | 67.7 | | | A:A | G:G |
| BS00023211 | 2D | 72.3 | 2D | 11.8 | G:G | A:A |
| BS00021950 | 2D | 84.2 | | | G:G | A:A |
| BS00062864 | 2D | 96.1 | | | T:T | C:C |

| | | | | | | |
|---|---|---|---|---|---|---|
| BS00082501 | 3D | 34.7 | | | A:A | G:G |
| BS00037276 | 3D | 88.6 | | | G:G | C:C |
| BS00147238 | 3D | 97.3 | | | A:A | G:G |
| BS00140559 | 3D | 97.8 | | | T:T | C:C |
| BS00103682 | 4D | 69.6 | | | T:T | C:C |
| BS00023046 | 4D | 70.2 | 4D | 12.5 | T:T | C:C |
| BS00023014 | 4D | 105.4 | 4D | 53.0 | A:A | G:G |
| BS00163273 | 5D | 73.8 | | | A:A | G:G |
| BS00145481 | 5D | 73.8 | | | T:T | G:G |
| BS00148826 | 5D | 104.4 | | | A:A | G:G |
| BS00064691 | 5D2 | 25.0 | | | G:G | T:T |
| BS00067650 | 5D2 | 43.2 | | | T:T | C:C |
| BS00055493 | 5D2 | 70.3 | | | T:T | G:G |
| BS00180355 | 5D2 | 70.9 | | | G:G | A:A |
| BS00084133 | 5D2 | 75.9 | 5D | 154.2 | C:C | T:T |
| BS00141290 | 5D2 | 78.1 | 5D | 160.5 | A:A | G:G |
| BS00021991 | 5D2 | 85.3 | 5D | 163.7 | A:A | G:G |
| BS00075107 | 5D2 | 94.2 | | | G:G | C:C |
| BS00182757 | 6D | 0.0 | 6D | 3.4 | T:T | C:C |
| BS00140915 | 6D | 3.4 | 6D | 1.2 | C:C | T:T |
| BS00042153 | 6D2 | 8.6 | | | A:A | C:C |
| BS00093153 | 6D2 | 9.2 | | | G:G | A:A |
| BS00117655 | 6D2 | 19.5 | | | C:C | C:T |
| BS00107889 | 6D2 | 51.1 | 6D | 115.7 | T:T | G:G |
| BS00181029 | 6D2 | 65.4 | 6D | 132.2 | G:G | A:A |
| BS00150885 | 6D2 | 70.5 | | | G:G | A:A |
| BS00139360 | 7D | 22.0 | | | T:T | G:G |
| BS00077071 | 7D | 37.1 | | | C:C | T:T |
| BS00023150 | 7D | 40.4 | | | C:C | T:T |
| BS00023045 | 7D | 115.8 | 7D | 122.2 | A:A | G:G |
| BS00108793 | 7D2 | 0.5 | | | G:G | T:T |
| BS00062860 | 7D2 | 4.0 | | | G:G | A:A |
| BS00166010 | | | 4D | 3.6 | T:T | C:C |
| BS00065168 | | | 4D | 53.0 | C:C | T:T |
| BS00130342 | | | 5D | 31.8 | A:A | G:G |
| BS00077450 | | | 5D | 38.0 | T:T | A:A |
| BS00074877 | | | 5D | 79.8 | G:G | C:C |
| BS00160381 | | | 6D | 77.2 | T:T | C:C |
| BS00105996 | | | 7D | 12.8 | A:A | T:T |
| BS00123834 | | | 7D | 42.2 | T:T | C:C |
| BS00064002 | | | | | T:T | C:C |

**Supplementary Table S5.** Selection of 33 A- and 33 B-genome KASP to for background selection in marker-assisted backcrossing. Markers were selected to be evenly distributed and co-dominant (where possible) and polymorphic between the recurrent parent Paragon and the AB-genome synthetic wheat donor Hoh-501. Chromosome allocation and genetic distance (cM) was based on the Avalon x Cadenza (AxC) population. All data available from https://www.cerealsdb.uk.net.

| SNP ID | AxC_Chr | AxC cM | SNP calls | |
|---|---|---|---|---|
| | | | Paragon | Hoh-501 |
| BS00022701 | 1A | 25.7 | T:T | C:C |
| BS00022934 | 1A | 33.2 | G:G | T:T |
| BS00078982 | 1A | 46.0 | A:A | C:C |
| BS00076668 | 1A | 55.6 | G:G | A:A |
| BS00039377 | 1A | 69.7 | C:C | A:A |
| BS00030767 | 1B | 28.8 | G:G | C:C |
| BS00099829 | 1B | 46.7 | T:T | G:G |
| BS00067434 | 1B | 76.7 | C:C | T:T |
| BS00032076 | 1B | 119.9 | A:A | C:C |
| BS00022487 | 2A | 40.4 | G:G | G:A |
| BS00001108 | 2A | 57.1 | G:G | C:C |
| BS00031140 | 2A | 63.5 | C:C | T:T |
| BS00089310 | 2A | 118.4 | T:T | A:A |
| BS00022209 | 2A | 128.2 | A:A | G:G |
| BS00081871 | 2B | 11.9 | C:C | A:A |
| BS00074661 | 2B | 72.8 | A:A | G:G |
| BS00023145 | 2B | 82.3 | A:A | T:T |
| BS00064740 | 2B | 95.6 | C:C | A:A |
| BS00046601 | 2B | 119.6 | C:C | T:T |
| BS00022016 | 3A | 64.6 | G:G | T:T |
| BS00039925 | 3A | 73.1 | G:G | T:T |
| BS00041462 | 3A | 91.1 | A:A | G:G |
| BS00088756 | 3A | 101.4 | T:T | C:C |
| BS00022735 | 3A | 167.0 | G:G | A:A |
| BS00089103 | 3B | 2.4 | T:T | C:C |
| BS00032912 | 3B | 61.7 | A:A | G:G |
| BS00070826 | 3B | 110.9 | C:C | G:G |
| BS00030651 | 3B | 182.3 | T:T | C:C |
| BS00022441 | 3B | 215.5 | A:A | G:G |
| BS00036493 | 4A | 8.8 | C:C | A:A |
| BS00049911 | 4A | 29.2 | G:G | T:T |
| BS00023220 | 4A | 85.8 | T:T | A:A |
| BS00021957 | 4A | 104.3 | A:A | G:G |
| BS00097249 | 4A | 113.9 | T:T | A:A |
| BS00044374 | 4B | 37.0 | A:A | C:C |
| BS00031139 | 4B | 52.8 | C:C | T:T |
| BS00037020 | 4B | 64.0 | G:G | T:T |
| BS00068539 | 4B | 99.2 | G:G | A:A |
| BS00022366 | 4B | 107.2 | G:G | A:A |
| BS00034303 | 5A | 1.1 | A:A | C:C |
| BS00022457 | 5A | 67.1 | C:C | T:T |

| | | | | |
|---|---|---|---|---|
| BS00105208 | 5A | 107.0 | G:G | C:C |
| BS00061064 | 5B | 43.7 | T:T | C:C |
| BS00022673 | 5B | 134.6 | T:T | C:C |
| BS00028197 | 5B | 180.4 | G:G | C:C |
| BS00036667 | 5B | 224.2 | T:T | A:A |
| BS00021982 | 6A | 8.9 | A:A | G:G |
| BS00031178 | 6A | 64.7 | C:C | T:T |
| BS00073872 | 6A | 101.8 | C:C | T:T |
| BS00021965 | 6A | 133.0 | G:G | A:A |
| BS00078124 | 6A | 157.4 | T:T | C:C |
| BS00030457 | 6B | 16.9 | A:A | G:G |
| BS00023208 | 6B | 40.8 | T:T | G:G |
| BS00023224 | 6B | 82.2 | G:G | C:C |
| BS00046264 | 6B | 104.2 | C:C | T:T |
| BS00110651 | 6B | 127.8 | C:C | T:T |
| BS00012880 | 7A | 23.9 | T:T | C:C |
| BS00023055 | 7A | 59.5 | A:A | G:G |
| BS00099804 | 7A | 100.9 | C:C | T:T |
| BS00031028 | 7A | 135.2 | A:A | G:G |
| BS00071736 | 7A | 217.3 | T:T | G:G |
| BS00070791 | 7B | 5.8 | T:T | C:C |
| BS00101408 | 7B | 9.8 | A:A | C:C |
| BS00023034 | 7B | 46.1 | T:T | C:C |
| BS00022195 | 7B | 60.7 | C:C | G:G |
| BS00050328 | 7B | 66.5 | G:G | A:A |

**Supplementary Table S6.** Graphical genotype selections from the populations across all three sub-genomes (A, B from NIAB_AB and D from NIAB_D). The individual and marker number in the selections is shown. The average length of selected introgressions is shown in Mb, as is the SNP marker spacing from these datasets. The average segment number refers to the average number of introgressions each line had per target genome. The percentage background recovery refers to the proportion of 'Paragon-like' SNPs in each line estimated using the array consensus map (Allen et al., 2017).

| Genome | No. Individuals | No. Markers | Av. int. length* | Av. segment no. | % Background recovered | Marker spacing (Mb) |
|---|---|---|---|---|---|---|
| A | 44 | 1444 | 102 | 2.3 | 96.0 | 3.4 |
| B | 33 | 1707 | 109 | 1.9 | 95.6 | 3.0 |
| D | 32 | 644 | 129 | 2.2 | 95.6 | 6.1 |

*Denotes average introgression length in Mb
